# Active site electrostatic effects modulate DBO inhibitor potency against OXA-48 variants

**DOI:** 10.64898/2026.11.04.764282

**Authors:** Daojiong Wang, Roop Chahal, Viivi H. A. Hirvonen, James Spencer, Marc W. van der Kamp

## Abstract

The OXA-48 class D *β*-lactamase and its variants are clinically concerning through their carbapenemase and cephalosporinase activity. Effective inhibitors are therefore urgently needed. Diazabicyclooctanes (DBOs) are newly introduced non *β*-lactam based inhibitors, but their potency is affected by mutations, with the more recently developed nacubactam consistently less potent than avibactam, the first DBO, towards OXA-48 variants. This difference is most pronounced for OXA-48 compared to its variants OXA-163 and -405, which contain mutations in the *β_5_–β_6_* loop and preferentially hydrolyse cephalosporins over carbapenems. Here we use multiscale simulations to uncover the origin of these differences after determining that the second step of DBO acylation is rate-limiting. Using QM/MM MD simulations we show that nacubactam exhibits a consistently higher energy barrier than avibactam for covalent inhibition of OXA-48-like enzymes, a difference attributable to DBO-specific changes in the active site electric field. Simulations of non-covalent complexes further indicate that weaker binding, due to electrostatic repulsion between the positively charged nacubactam tail and Arg214 in the *β_5_–β_6_* loop, partially explains the lower potency of nacubactam towards OXA-48, compared to OXA-163 and -405. Combining computational predictions of covalent reaction rates and non-covalent binding affinities accurately reproduces the relative order of experimental IC_50_ values, and predicts potency towards OXA-517, a recently discovered OXA-48 variant capable of hydrolysing many antibiotics. This work provides mechanistic insight into the inhibition of class D *β*-lactamases and reveals the origins of differential DBO potency towards OXA-48 variants, highlighting how subtle differences between enzymes modulate inhibitory effects.

## Introduction

The *β*-lactam class of antibiotics are the most widely used antimicrobial agents for treating bacterial infections around the world ^1, 2^. However, the effectiveness of *β*-lactam antibiotics is rapidly declining due to the emergence of antibiotic resistance, mainly driven by *β*-lactamases (BLs), enzymes produced by bacteria that hydrolyse the *β*-lactam functional group, leading to antibiotic inactivation ^3^. According to the Ambler classification system, BLs can be divided into four classes: A, B, C, and D ^4^. Classes A, C and D share a conserved serine residue in the active site, which performs nucleophilic attack on the *β*-lactam ring, while class B BLs are zinc-dependent, and hydrolyse *β*-lactams through a distinct, metal-mediated mechanism ^5^. Therefore, classes A, C and D BLs are referred to as serine-*β*-lactamases (SBLs), and class B BLs are metallo-*β*-lactamases (MBLs). Among SBLs, class D enzymes have increasingly attracted attention because many of their members are capable of hydrolysing carbapenems, which were once considered ‘last-resort’ antibiotics ^6^. Hence, the development of new strategies to combat antibiotic resistance is urgently needed.

One of the most effective strategies to restore antibiotic efficacy is combining antibiotics with *β*-lactamase inhibitors. *β*-lactamase inhibitors in clinical use can be divided into three classes according to their structural features: *β*-lactam based inhibitors, diazabicyclooctane (DBO) based inhibitors and boronic acid based inhibitors ^7^. DBOs exhibit a wider inhibitory spectrum (including many class A, C and D BLs) than many other inhibitors, and their reversible covalent inhibition mechanism via recyclization contributes to improved efficacy ^5^. The DBO avibactam was the first clinically approved non-*β*-lactam *β*-lactamase inhibitor, used in combination with ceftazidime for treating resistant Gram-negative bacterial infections ^8, 9^. More recently (2024), avibactam was approved in combination with aztreonam for treating serious Gram-negative bacterial infections ^10^. Several other DBO inhibitors combined with carbapenems have been approved or entered clinical trials. Relebactam was approved in combination with imipenem/cilastatin in 2019 ^11^. Another DBO inhibitor, nacubactam, has entered clinical trials in combination with meropenem ^12^. Nacubactam and zidebactam also show inhibition of bacterial Penicillin-binding protein 2 (PBP2) which is essential for cell wall synthesis^13, 14^. Like *β*-lactam based inhibitors, DBOs inhibit SBLs by reacting with the active site serine nucleophile to form long-lived covalent complexes, but unlike *β*-lactams, such inhibition is reversible: BL-bound avibactam is found to recyclize to the ring-closed intact form in many class A and C BLs ^15^. However, the recyclization rate of avibactam in many class D BLs was found to be extremely slow, thereby resulting in an “almost irreversible” inhibition ^16, 17^.

Compared with other SBLs, class D BLs uniquely possess a carboxylated Lys73 (KCX) in the active site, which functions as a general base during *β*-lactam hydrolysis by activating the serine nucleophile for acylation, and a water molecule for deacylation ^6, 18^. Carboxylation of Lys73 has been reported as essential for antibiotic hydrolysis ^19, 20^. Recently, several computational studies have demonstrated that KCX also plays a crucial role in acylation with avibactam ^21, 22^. The mechanism of DBO acylation in OXA-48, a member of class D BLs, consists of two steps (**Figure 1**). After the inhibitor forms a Michaelis complex (MC) with OXA-48, KCX acts as a general base to activate Ser70 via a proton transfer. The deprotonated Ser70 then performs a nucleophilic attack on the electrophilic DBO carbonyl carbon, forming a covalent bond, and leading to a negatively charged tetrahedral intermediate (TI). This step is similar to *β*-lactam antibiotic acylation of OXA-48, and the hydrogen bonds between the negatively charged oxygen and the oxyanion hole residues, which stabilise the tetrahedral intermediate (TI), are conserved ^20, 23^. Subsequently, the second step involves breakage of the DBO γ-lactam ring, along with a proton transfer from Lys208 to the (former) γ-lactam N via Ser118, resulting in formation of a stable acylenzyme complex (AC). Unlike *β*-lactam hydrolysis, the DBO acylenzyme does not undergo deacylation to release the hydrolysed product, and thereby inhibits the enzyme.

**Figure 1.**
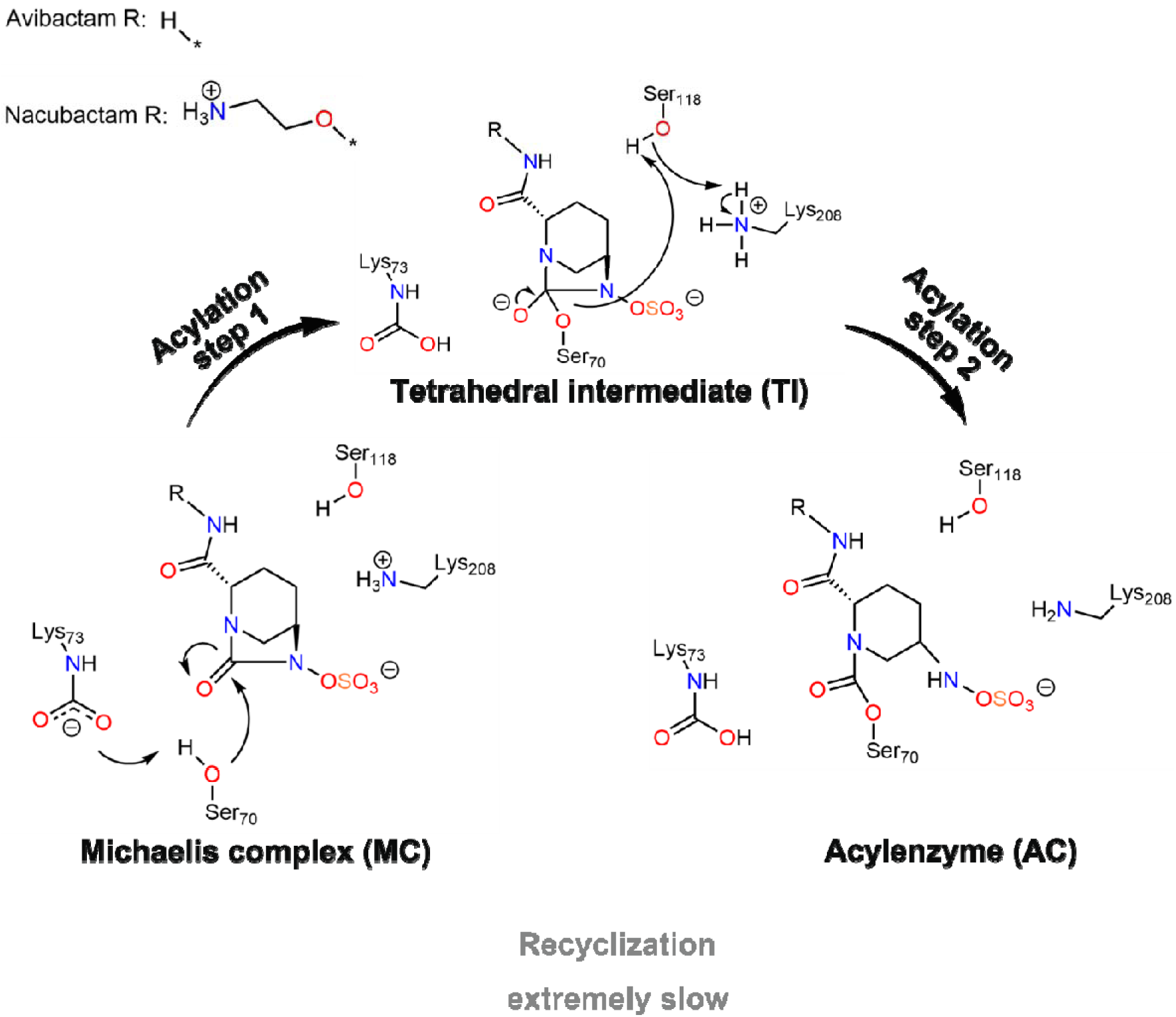
Reaction mechanism of DBO acylation of OXA-48 ^21, 22^.

OXA-48-like enzymes are a clinically relevant family within the class D BLs, that can confer resistance to an extended range of *β*-lactams ^18^ and that contains multiple variants with several members being wide-spread. Sequence variation at two regions located near the active site, not directly involved in the reaction, can impact activity of these enzymes. The flexibility of the Ω-loop (OXA-48 residues Tyr144 to Arg163) has been reported to correlate with the functional trade-off between carbapenem- and cephalosporin-hydrolysing activities in OXA-48-like enzymes ^24, 25^. The other region is the *β_5_*–*β_6_* loop (residues Thr213 to Lys218 in OXA-48), which can form a salt bridge interaction with the Ω-loop. Mutations that impair this interaction lead to a less stable Ω-loop and also introduce a trade-off between carbapenemase and cephalosporinase activities ^26^. Two natural OXA-48 *β_5_*–*β_6_* loop variants, OXA-163 and -405 (**Figure 2**), have been reported to exhibit extended activity towards cephalosporins but reduced carbapenemase activity ^27, 28^. By contrast, OXA-517 was reported to hydrolyse both cephalosporins and carbapenems ^29^.

**Figure 2.**
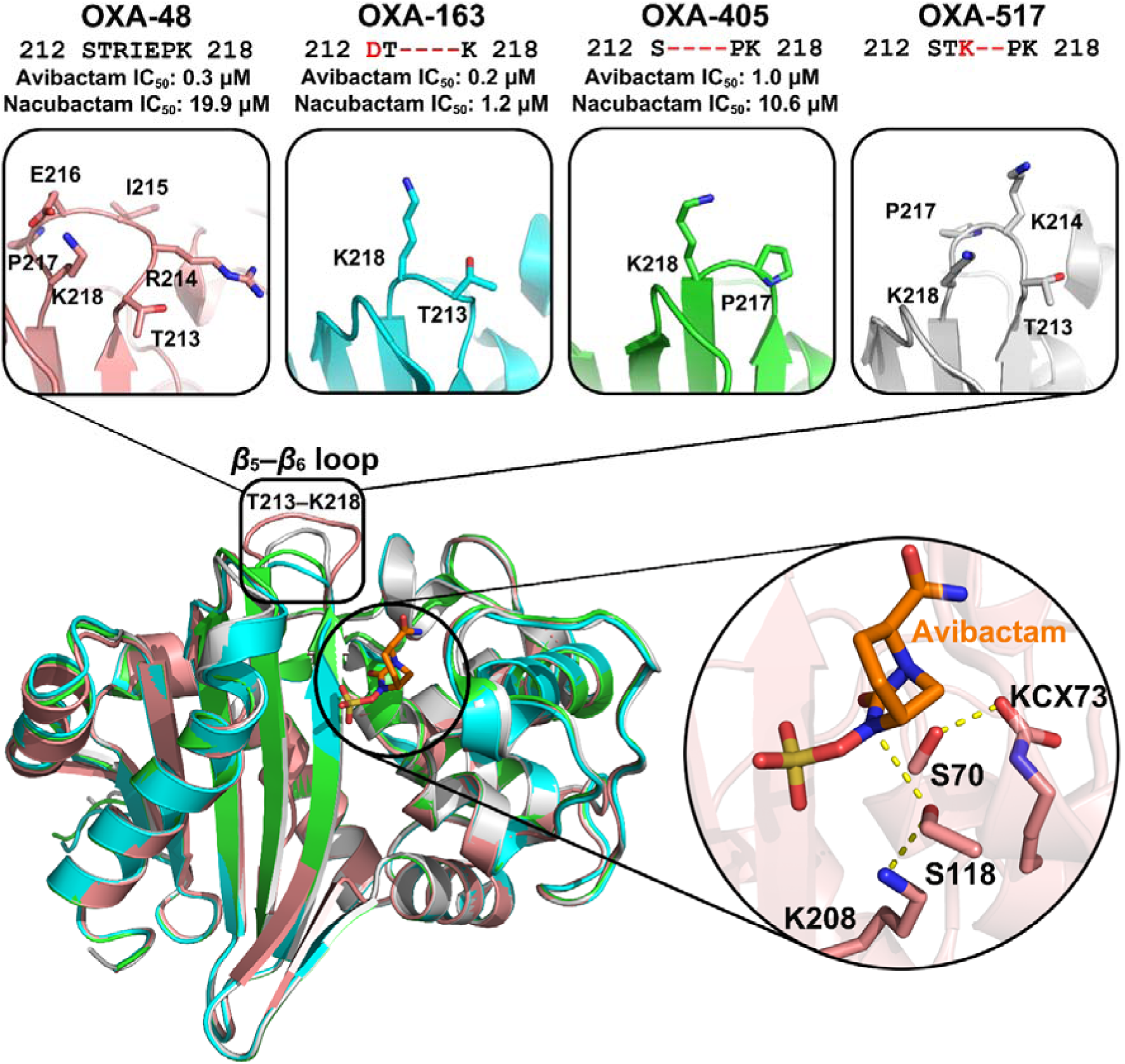
Aligned crystal structures of OXA-48 (pink, PDB ID: 4S2K ^31^, chain A, avibactam acylenzyme complex), OXA-163 (cyan, PDB ID: 9H14 ^30^, chain A, avibactam acylenzyme complex), OXA-405 (green, PDB ID: 9H17 ^30^, chain A, avibactam acylenzyme complex) and OXA-517 (grey, PDB ID: 6HB8 ^29^, chain A, apo). For clarity, the free avibactam derived from the acylenzyme crystal structure is shown as sticks (orange) only in OXA-48. The positions of the avibactam and nacubactam cores are highly similar. Structural differences in the *β_5_*–*β_6_*loop are highlighted in the inserts, together with sequences of the *β_5_*–*β_6_* loop and experimental IC_50_ values from a published study ^30^. The key hydrogen bonds involved in proton transfer are also highlighted in the inset (yellow dashed lines), where avibactam (orange) and the sidechains of key residues (pink) are shown as sticks.

Recent experimental data reveal distinct inhibitory effects of avibactam and nacubactam against OXA-48 variants ^30^. Both avibactam and nacubactam share very similar chemical structures (**Figure 1**). Nevertheless, nacubactam exhibits consistently higher IC_50_ values against OXA-48 variants compared with avibactam (**Figure 2**). Notably, for OXA-48, the IC_50_ value of nacubactam is much higher than that of avibactam, whereas the difference is much smaller for OXA-163 and -405. The only difference among these OXA-48 variants lies in the *β_5_*–*β_6_* loop and its surrounding residues (**Figure 2**). This suggests that the nature of *β_5_*–*β_6_* loop is important not only for antibiotic hydrolysis but also for DBO inhibition.

Here, we use detailed, multiscale simulations to understand the precise molecular basis of the differences in DBO inhibition resulting from *β_5_*–*β_6_* loop mutations. After establishing the detailed mechanism of DBO acylation, reaction simulations of the rate-limiting step in all variants reveal a trend consistent with experimental IC_50_ values ^30^: nacubactam always exhibits higher free energy barriers than avibactam, due to small changes in the active site electric field. Molecular dynamics simulations of non-covalent Michaelis complexes further show that (in-tact) nacubactam binds more weakly than avibactam, especially in OXA-48. By considering both the predicted free energy barriers and (relative) binding energies, the order of the experimentally obtained IC_50_ values in OXA-48, -163 and -405 can be accurately reproduced. We then use the same simulations to estimate IC_50_ values of avibactam and nacubactam for OXA-517. Taken together, our results reveal the basis for differential inhibitory effects of different DBOs on class D *β*-lactamases in atomic detail, providing information that can underpin rational inhibitor design against this family of clinically important *β*-lactamases.

## Methods

### Simulation system setup and preparation

The crystal structures of DBO-OXA-48, -163, and -405 acylenzyme complexes were obtained from the PDB database (for avibactam, PDB IDs 4S2K ^31^, 9H14 ^30^, and 9H17 ^30^, respectively, and for nacubactam, 9H12 ^30^, 9HPV ^30^, and 9H18 ^30^). In all cases, chain A from the crystal structures was used. The Michaelis complexes of DBOs binding with OXA-48, - 163 and -405 were generated from their corresponding acylenzyme complex structures by breaking the bond between Ser70 and DBOs and reforming the diazabicyclooctane ring in Pymol 3.1.3, followed by relaxation (see next section). The Michaelis complexes of DBOs binding with OXA-517 were constructed by modelling avibactam and nacubactam into the apo form crystal structure of OXA-517 (PDB ID: 6HB8 ^29^, chain A) through alignment with the avibactam-OXA-48 and nacubactam-OXA-48 Michaelis complexes respectively. The crystal waters were retained and residues corresponding to the signal peptide (residue number 1 to 22) were removed. The missing atoms of the carboxylated Lys73 (KCX) were added manually in the avibactam-OXA-48, nacubactam-OXA-48, nacubactam-OXA-405 complexes. The partial charges of free DBOs were calculated using the AM1-BCC model ^32^ with antechamber in AmberTools24 ^33^. The atom types of bonded parameters of DBOs were described using the GAFF2 force field ^34^, and proteins were treated with the Amber ff14SB force field ^35^. The protonation states of ionizable residues and DBOs at pH 7.0 were determined by using PropKa3.1 ^36^, and the histidine tautomers were predicted by the reduce program in AmberTools24. To minimise the impact of protonation state differences on the results, one consistent set was used: all residues were in their standard protonation states (Asp and Glu deprotonated; Lys and Arg protonated), with all His residues being neutral and singly protonated on NE2 except His140 singly protonated on ND1. Avibactam has a net charge of −1e, while nacubactam is overall neutral, with the tail amino group protonated ^30^. The Michaelis complexes were solvated in a rectangular box of TIP3P water ^37^, with a minimum distance between the protein and the box edge of 10 Å. The whole systems with solvent water box were neutralized by replacing random bulk water molecules with Na^+^ ions. Three Na^+^ ions were added to the avibactam-OXA-48, -163 systems, and two Na^+^ ions were added to the nacubactam-OXA-48, -163, and avibactam-OXA-405 systems, one Na^+^ ion was added to the nacubactam-OXA-405 and avibactam-OXA-517 systems, no ions were added to the nacubactam-OXA-517.

### Michaelis complex MM MD simulation and analysis

All systems were minimised in three stages: minimising only the solvent while applying positional restraints on the rest of the system; then restraining Cα atoms only and minimising the remaining atoms; finally minimising the whole system without positional restraints. All minimisation steps have 1000 cycles with steepest descent followed by another 1000 cycles with conjugate gradient, and the restraint force constant was set to 50 kcal mol^−1^ Å^−2^. After minimisation, the system was heated from 50 to 300 K with pressure of 1 bar over a period of 50 ps. Langevin dynamics were used with a collision frequency of 0.2 ps^−1^. The Berendsen barostat was used with isotropic position scaling. During all equilibration and production simulations, the systems were simulated in the NPT ensemble at 300 K and 1 bar. Periodic boundary conditions were applied and the SHAKE algorithm was applied to fix all bond lengths involving hydrogen atoms. A timestep of 2 fs was used and the cutoff radius for non-bonded interactions was set to 8.0 Å. All simulations were conducted using Amber24 ^33^ and trajectories were analysed by using CPPTRAJ ^38^. For hydrogen bond analysis, a hydrogen donor-acceptor distance cutoff of 3.5 Å, and a donor-hydrogen-acceptor angle between 135° and 180° were used.

Three sets of production simulations were conducted with different harmonic restraints applied for different purposes. The force constant for all restraints is 10 kcal mol^−1^ Å^−2^ or 10 kcal mol^−1^ rad^−2^. (i) Five independent 20 ns equilibration simulations were conducted for all systems followed by 100 ns production. The trajectories were saved every 10 ps, resulting in 10000 frames per replica for analysis. A one-sided harmonic restraint was applied to maintain the binding of the DBOs to the active site by keeping the distance between the DBO carbonyl oxygen and the backbone amide hydrogen of Ser70 within 2.4 Å. A two-sided harmonic restraint was also applied on Ser70 χ1, allowing restraint-free sampling from 270° to 320° to ensure the hydroxyl group pointed to the KCX. These production simulations were used to analyse conformations and interactions, and provide initial structures for subsequent umbrella sampling simulations. (ii) One 10 ns production simulation was performed for avibactam-OXA-48 and -163 systems, and the trajectories were saved every 5 ps, yielding a total of 2000 frames per system. Four restraints were applied to ensure the binary complex is reactive for acylation. Three one-sided distance harmonic restraints came into force when the distances exceeded 2.0 Å : between the avibactam carbonyl oxygen and the backbone amide hydrogen of Ser70/Tyr211, and between KCX@OQ1 and Ser70@HG. A two-sided dihedral angle harmonic restraint was also applied on Ser118 χ1 with restraint-free sampling from 190° to 230°. The final frames of the trajectories were used for QM/MM potential energy calculations described below. (iii) Ten independent 10 ns production simulations were conducted for all systems. The trajectories were saved every 10 ps, resulting in 1000 frames per replica for subsequent MM-PBSA binding free energy calculations. The same harmonic restraints as those used during the 20 ns equilibration of the 100 ns MM MD simulations of Michaelis complexes described above were applied in both heating and production stages.

### QM/MM reaction mechanism calculations

The last frames of the 10 ns MM MD simulation of the avibactam-OXA-48 and -163 systems were used as starting structures for 100 ps QM/MM MD simulation (for structural relaxation). The QM region was described using the semi-empirical method DFTB2 (SCC-DFTB) ^39^. The whole of avibactam and part of the sidechains of Ser70, KCX, Ser118 and Lys208 are included in the QM region **(Figure S1)**. For the avibactam-bound systems, each QM region contains 56 atoms (including 4 link atoms) with a total charge of −1 e, whereas the nacubactam-bound systems comprise 66 atoms (including 4 link atoms) and is neutral overall. The same three distance restraints as in the previous MM MD simulations were applied. An additional one-sided harmonic restraint (force constant also set to 10 kcal mol^−1^ Å^−2^) was applied to maintain the distance between Ser70@OG and its hydrogen (Ser70@HG) below 1.0 Å. Simulation settings were identical to the previous MM simulations, apart from using a 1 fs timestep and 10.0 Å non-bonded interactions cutoff (and no SHAKE applied on the QM region).

Two frames per system, corresponding to different hydration states around the avibactam moiety (**Figure S2**), were extracted from the 100 ps QM/MM MD simulation. The structures were stripped of any residues with any atoms further than 17 Å from the avibactam carbonyl oxygen. Atoms which are closer than 10 Å to the carbonyl oxygen were included in the active region and allowed to move during subsequent potential energy calculations. The force field parameters for MM atoms were taken from the prior MM MD simulations. The CI-NEB (climbing image nudged elastic band) ^40^ implemented in ORCA 6.0 ^41^ was used to investigate the minimum energy path of DBO acylation of class D *β*-lactamases, and identify the rate-limiting step.

An optimized starting structure for the TI (tetrahedral intermediate) was generated from the stripped system by applying a constraint to change the bond arrangement (distance between Ser70@HG and KCX@OQ1 set to 1.0 Å). The structure was first minimised using GFN2-xTB (XTB2) ^42^, with a subsequent full free minimisation using r^2^SCAN-3c ^43^. The starting structures of MC (Michaelis complex) and AC (acylenzyme) were obtained from the optimized TI structure directly using r^2^SCAN-3c, first by applying constraints, followed by full free energy minimisation. For MC generation, the distance between Ser70@OG and Ser70@HG was set to 1.0 Å, whereas for AC, the distances between Ser70@HG and KCX@OQ1, Ser118@HG and the γ-lactam ring nitrogen of avibactam, Ser118@OG and the reactive Lys208@HZ were set to 1.0 Å. The RMS gradient tolerance was set to be below 5×10^−5^ a.u. during all optimizations. The reaction pathways were separated into two steps: the TI formation from the MC and the AC formation from the TI. The pre-optimized starting structures were used for CI-NEB calculations with frozen ends at the r^2^SCAN-3c level. The images were set to be redistributed along the path every 50 iterations. The default convergence criteria were kept during the CI-NEB calculations. Then the single point energy calculations of converged images were performed on ωB97X/def2-TZVP level ^44, 45^. The CI-NEB calculations were repeated with water molecules forming H-bonds with the avibactam carbonyl oxygen and the reactive oxygen of KCX included in the QM region (**Figure S2**). CI-NEB calculations with free ends, using r^2^SCAN-3c optimized starting structures, were performed at the ωB97X/def2-TZVP level of theory to validate the reliability of the reaction pathway obtained from r^2^SCAN-3c. The RIJCOSX approximation with the default auxiliary def2/J basis set were applied ^46, 47^. The L-BFGS optimizer ^48^ was used for all geometry optimizations and CI-NEB calculations.

### QM/MM MD reaction simulation and analysis

After identifying the rate-limiting step of DBO acylation (from TI to AC), 2D QM/MM US (umbrella sampling) MD was performed to investigate the inhibition effect differences of avibactam and nacubactam against OXA-48, -163, -405 and -517. In order to minimize the computational cost, the semi-empirical method XTB2, which correlates the best with high-level QM method, was used to describe the QM region (**Figure S3**) (see Supplementary Note S1 for details). One frame per replica was extracted from each of the five independent 100 ns Michaelis complex MM MD simulation, in which a strong hydrogen bond is formed between KCX@OQ1 and Ser70@OG, and the γ-lactam ring nitrogen is pointed towards Ser118 (positioned for a stereo-electronically favoured reaction) ^49^. The force constant for all restraints and reaction coordinates used in QM/MM US MD simulations described below was set to 100 kcal mol^−1^ Å^−2^ or 100 kcal mol^−1^ rad^−2^.

First, a 50 ps QM/MM MD simulation was conducted to relax the structure. Five replicas were performed for each system. The same QM region mentioned above was set (**Figure S1**). To preserve the reactive conformation of Michaelis complex, restraints similar to those used in the previous 100 ps QM/MM MD relaxation in QM/MM potential energy calculations were applied, and several new restraints were additionally introduced (see Supplementary Note S2). Simulation settings were identical to the previous 100 ps QM/MM MD relaxation mentioned above.

For the following QM/MM US MD simulations, simulation settings were identical to those of the previous 100 ps QM/MM MD relaxation, except that the non-bonded interaction cutoff was set to 8.0 Å. The last frame of the simulation was used to generate the TI structure using 1D QM/MM US MD. A single combined reaction coordinate was employed: distance (−*d*[KCX@OQ1, Ser70@HG] − *d*[Ser70@OG, DBO@C]). The reaction coordinate changes from −4.2 Å to −2.5 Å over 18 US windows in total. Each window has 2 ps sampling time with 1 fs timestep. Same restraints as those used in the previous 50 ps QM/MM MD simulation were applied, except for those involved in the reaction coordinate. In addition, three one-sided harmonic restraints were applied on the NH_3_^+^ group of Lys208 to maintain N-H bond lengths below 1.0 Å and to prevent any proton transfer between the NH_3_^+^ group and the DBO sulfate ester group.

The generated TI structure was then used as the starting point for simulating the second step through 2D QM/MM US MD simulation. One reaction coordinate is the proton transfer coordinate among DBO, Ser118 and Lys208: distance (−*d*[DBO@N, Ser118@HG] + *d*[Ser118@HG, Ser118@OG] − *d*[Ser118@OG, Lys208@HZ] + *d*[Lys208@HZ, Lys208@NZ]) which shows the breakage of O-H bond in Ser118 and N-H bond in Lys208, accompanied by the formation of O-H bond in Ser118 and N-H bond in the DBO. The other is the C-N bond breakage in the DBO (*d*[DBO@N, DBO@C]). From the TI to AC, the proton transfer coordinate changes from −2.2 Å to 2.0 Å, while for C-N bond cleavage, it changes from 1.6 Å to 2.7 Å. The step size was set to 0.1 Å. Fewer restraints were applied in this step: only the two-sided dihedral angle restraint used to maintain the DBO in chair conformation, and those one-sided distance restraints used to prevent proton transfer from Ser118 or Lys208 to the DBO sulfate ester group. Each US window was equilibrated for 2 ps first, followed by 20 ps of production sampling with a 1 fs timestep. 516 US windows were used in total for each replica. The US calculations were performed by Amber24 and reaction coordinate values were saved every step and analysed using the Weighted Histogram Analysis Method (WHAM) ^50^. FES (free energy surface) was generated and plotted by the Minimum Energy Path Surface Analysis (MEPSA) program ^51^. Frames from the US windows where the transition state is located were used for electric field calculations using FieldTools ^52^. The effective electric field was calculated along the DBO C-N bond. Atoms directly involved in the reaction, their covalently bonded neighbours (i.e., atoms directly attached to the C or N atoms of the DBO, including their hydrogens), as well as the sidechains and Cα atoms of Ser70, Ser118, and Lys208 were excluded from the electric field calculations. For each simulated system, 100 frames were used (20 frames per transition state window, 5 replicas).

The same generated TI structures were used for simulating the first step reaction of avibactam and nacubactam with OXA-48 and -163 through 2D QM/MM US MD simulation as well. The simulation settings and analysis methods remained unchanged, except that the QM region was described using RM1 ^53^, which showed the best agreement with high-level QM energies for geometries along the first step reaction (**Figure S4**). The detailed information about reaction coordinates and restraints applied during the simulation can be found in Supplementary Note S3.

### Binding energy (MM-PBSA) and relative binding free energy calculation

MM-PBSA (with energy decomposition per residue) was performed to calculate the binding energy of DBOs to the different enzyme variants using MMPBSA.py ^54^ from AmberTools24. For each system, ten independent trajectories from 10 ns MM MD simulations of Michaelis complexes were used. The alchemical transfer method (ATM) ^55, 56^ implemented in BioSimSpace ^57^ with the OpenMM ^58^ engine, was used to calculate the relative binding free energy between different DBOs binding with the same enzyme variant. Nacubactam was set as bound ligand at the beginning. Hence, final frames from 10 ns MM MD simulations for both the OXA-48 and OXA-163 nacubactam Michealis complexes were extracted as the initial structures for the calculation. The starting structure of avibactam was generated by removing the tail of nacubactam and replacing the tail oxygen with hydrogen. Further details of the ATM calculations can be found in Supplementary Note S4.

## Results and Discussion

### Potential energy profiles reveal the second step of DBO acylation as rate-limiting

To elucidate the details of the DBO acylation reaction and understand the origin of differential inhibitory effects of OXA-48 variants through MD simulations, the rate-limiting step of the reaction needs to be identified. Two different hydration states around the DBO moiety were observed in MM MD and QM/MM MD simulations of the non-covalent (Michaelis) complexes of OXA-48, -163, -405 and -517, with either avibactam or nacubactam (**Figure S2**). Hydration state I harbours an additional water molecule in the active site, which can form a H-bond to the carbonyl oxygen of while in state II, no such solvent H-bond interaction is present. Both hydration states are frequently sampled in the 100 ns MM MD simulations of the Michaelis complexes (**Table S1**). Approximately 1/3 of the snapshots corresponded to hydration state I, except for OXA-405 with nacubactam, for which only 15.5% of frames adopt this state.

To establish the detailed DBO acylation reaction mechanism, DFT-level QM/MM potential energy calculations were performed for both hydration states for avibactam acylation reactions in OXA-48 and OXA-163 as representative systems. The climbing image nudged elastic band (CI-NEB) method was employed (at the r^2^SCAN-3c/ff14SB level), using representative enzyme-substrate conformations as starting points (see Methods). In OXA-48, different hydration states only have a minor impact on the structures along the minimum energy path (**Figure S5**). In the first step of acylation (Michaelis complex → tetrahedral intermediate), the potential energy starts to increase as the nucleophilic attack initiates. The proton transfer between KCX and Ser70 happens concerted with the nucleophilic attack, with the proton positioned nearly equidistant between these two residues just before the transition state (TS1) is reached. In the second step of acylation (tetrahedral intermediate → acylenzyme), the initial rise in energy and the transition state (TS2) is characterized by the proton transfer between S118 and avibactam, along with the C-N bond breakage of avibactam. The proton transfer between Lys208 and Ser118 occurs slightly later, revealing the order of the two proton transfer events.

The different hydration states do lead to a change in the relative contributions to the overall acylation barrier from the two reaction steps. When the initial complex is at hydration state II, the energy barrier of the first step increases, while that of the second step decreases (**Figure 3**), compared to hydration state I. A significant energy difference between the MC and TI was observed in hydration state II (12.7 kcal mol^−1^), while the difference is only 5.8 kcal mol^−1^ in hydration state I, suggesting that the additional water molecule stabilises the TI. By comparing the two transition states between hydration states I and II, we found that the locations of the second step transition states (TS2) are nearly identical. However, for the first step, the transition state (TS1) occurs slightly later in hydration state II, as indicated by the longer C-O bond distance (1.9 Å in hydration state I vs. 1.7 Å in hydration state II) (**Figure 3**).

**Figure 3.**
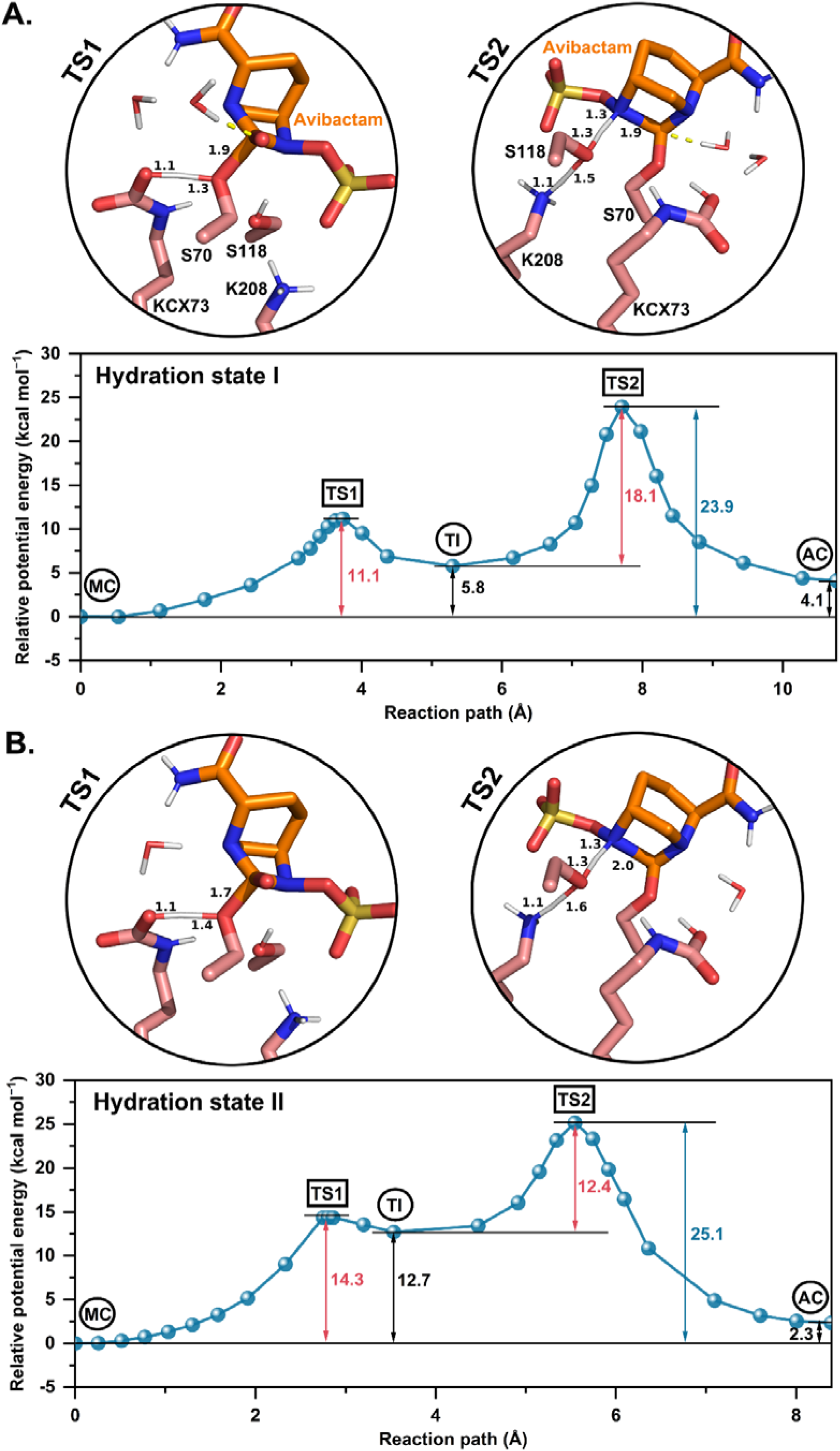
Potential energy profiles of avibactam acylation of OXA-48 in hydration states I (A) and II (B), with MC = Michaelis complex, AC = acylenzyme, TI = tetrahedral intermediate, TS = transition state. CI-NEB calculations were performed at the r^2^SCAN-3c/ff14SB level, with potential energies refined to the ωB97X/def2-TZVP//ff14SB level through single point energy calculations. The relative energies were determined using MC as the reference. The potential energy barriers for both steps are shown in red, and the overall barrier in blue, while the energies of TI and AC relative to MC are shown in black. The structures of the transition states (climbing images) are shown, with reactive residue sidechains (pink carbons) and avibactam (orange carbons) represented as sticks. Distances of changing bonds are labelled in Å. The solvent H-bond to avibactam carbonyl oxygen is indicated by a yellow dashed line.

As a relatively recent composite method with empirically fitted parameters, the reliability of r^2^SCAN-3c for geometry optimisation was validated by comparing the results of the avibactam acylation reaction in hydration state I of OXA-48 with those obtained from the CI-NEB calculations using ωB97X/def2-TZVP to describe the QM region. The potential energy barrier of the first step calculated by ωB97X/def2-TZVP//ff14SB CI-NEB is 11.6 kcal mol^−1^, which differs by only 0.5 kcal mol^−1^ from that obtained from the geometries at r^2^SCAN-3c/ff14SB level with ωB97X/def2-TZVP//ff14SB energy corrections (**Figure S6**). While the difference in the second step is slightly larger, the potential energy barriers are 18.2 and 16.5 kcal mol^−1^ for geometries generated at the r^2^SCAN-3c/ff14SB level with ωB97X/def2-TZVP//ff14SB energy corrections and at the ωB97X/def2-TZVP//ff14SB level, respectively. Notably, the geometries of the TS and minimum energy paths in both steps are similar (especially the first step) across the two QM levels. Our results show that r^2^SCAN-3c can achieve similar levels of conformational accuracy to those obtained using higher levels of QM theory, with significantly reduced computational cost ^43^.

Similar mechanisms and transition states were observed for reaction of OXA-163 with avibactam, indicating that OXA-48-like class D BLs share a common mechanism of reaction with DBOs, regardless of variant (**Figures S7** and **S8**). In OXA-163, hydration state I again stabilises the TI, resulting in a lower energy barrier for the first step compared with hydration state II, and a higher barrier for the second step. However, the energy barrier difference between the first and second step in hydration state I is only 1.6 kcal mol^−1^ (**Figure S7A**), much smaller than that in OXA-48 (7.0 kcal mol^−1^). Unlike OXA-48, in OXA-163 hydration state II, the second step energy barrier is still slightly higher than the first step (0.4 kcal mol^−1^) (**Figure S7B**). Both CI-NEB calculations of OXA-48 and -163 with avibactam show that the presence of the additional water molecule in hydration state I can help stabilise the TI. The carbonyl oxygen of avibactam engages in hydrogen bonding with the additional water molecule and also interacts with the backbone amides of Ser118 and Tyr211 in the oxyanion hole. Those two oxyanion hole interactions, plus the H-bond formed with the additional water, stabilise the negative charge on the (former) carbonyl oxygen of the tetrahedral intermediate ^23^. Although the stabilisation of the TI differs, the overall barrier for acylation is similar for both hydration states (**Figures 3** and **S7**).

The non-polarisable water model TIP3P, which was used for all water molecules, may lead to inaccurate prediction of electrostatic properties ^59, 60^. In order to better describe the interactions with the reacting atoms, the QM region was expanded to include the water molecules that form H-bonds with the carbonyl oxygen of avibactam and the KCX reactive oxygen (**Figure S2**). In both OXA-48 and -163, the energy differences between the MC and TI in hydration state I (which contains an additional water molecule in the active site) increase from 5.8 to 8.0 and from 8.7 to 10.5 kcal mol^−1^, and the respective energy barriers increase from 11.1 to 14.0 and from 12.4 to 14.0 kcal mol^−1^ (**Figures 3** and **S7** with water molecules not included in the QM region, **Figures S9** and **S10** with water molecules included in the QM region). Our results show that the stabilising effect of the additional water when described using TIP3P is overestimated, likely due to overpolarisation of the QM region by the point charges in the TIP3P model ^61^. Notably, even when both water molecules in hydration state I are included in the QM region (**Figure S2**), the second step is still predicted to have a higher barrier in both enzymes (**Figures S9** and **S10**). Although the overall barrier for acylation is slightly lower for hydration state I in all energy profiles, this is possibly compensated by this hydration state being less frequently sampled in our MD simulations (**Table S1**). Overall, although different hydration states affect the magnitude of the energy barriers associated with the individual steps and the stability of the TI, the TI remains a transient species rather than a long-lived state. Under most conditions the major energetic barrier associated with acylation is that for the second step (TI → AC), suggesting that the second acylation step is rate-limiting.

To compare our calculated barriers to experiment is not straightforward, as acylation rates are not measured directly. However, assuming *K*_i app_ is close to the dissociation constant for the non-covalent Michaelis complex *K*, the acylation rate *k*_2_ can be estimated from reported association rates *k*_2_/*K* ^16, 62, 63^. For OXA-48 with avibactam, thus estimated acylation rates from ref. ^62^ and refs. ^16, 63^ can be converted to reaction barriers (using the Eyring equation with 300 K) of 21.0 and 19.5 kcal mol^−1^, respectively. The most accurate overall barrier calculated here (CI-NEB calculations at the ωB97X/def2-TZVP//ff14SB level, **Figure S6**), 22.3 kcal mol^−1^, is consistent with this. Thakkur *et al*. applied metadynamics simulations using the PBE exchange-correlation functional and a plane wave basis set to study avibactam acylation of OXA-48 ^21^. They force the reaction directly from MC to AC, with the highest energy is reached between the TI and AC, consistent with our findings. There is only a very shallow minimum for the TI (similar to our results for hydration state II). By contrast, Lizana *et al*. explored the potential energy surface of acylation of OXA-24 by the DBO durlobactam using QM/MM (using the hybrid DFT with M06-2X/6-31+G(d,p)) and reported a relative energy of the TI that is, surprisingly, much lower than the MC (implying the TI to be a highly stable state) ^22^. Although this indicates that different methods and approaches can lead to different results regarding the detailed reaction energy profile, the consensus is that the highest energy barrier for avibactam acylation is associated with the second reaction step, as we show here for both OXA-48 and OXA-163 (independent of hydration state).

### Nacubactam exhibits a higher energy barrier due to changes in the electric field

To investigate differences in the free energy barriers for acylation of OXA-48 and -163 by avibactam and nacubactam, QM/MM umbrella sampling (US) simulations were performed. These simulations were started from the TI complexes, selecting GFN2-xTB (XTB2) ^42^ for the rate-determining second step and RM1 ^53^ for the first step as efficient QM methods, selected for optimal agreement with ωB97X/def2-TZVP energies (**Figures S3** and **S4**). We observed that the additional water molecule present in hydration state I quickly drifted away along the minimum free energy paths, and there was no correlation between the average number of solvent H-bonds with the former DBO carbonyl oxygen and the barrier of the second step (**Table S2**). Therefore, the final frame from each independent TI generation trajectory was used for US simulations, independent of the hydration state. Analysis of the free energy barriers for the different systems (based on Boltzmann-weighted averaging of obtained potential of mean force profiles) indicates that there is no significant difference between avibactam and nacubactam for the first step of acylation (**Figure S11**, **Table S3**), whereas a significant difference does arise in the second step, for both OXA-48 and OXA-163 (**Figure 4A, Table S3**). This difference results in an increase of the free energy barrier for acylation for nacubactam of approximately 4 kcal mol^−1^ compared to avibactam. Experimental data show that the IC_50_ values of nacubactam are significantly higher than those of avibactam (**Figure 2**) ^30^; our simulation results indicate that this difference is (at least in part) the results of changes in the efficiency of the second step of acylation, where nacubactam has a larger free energy barrier. Therefore, this second step of the acylation reaction was investigated further, to understand the origin of this difference.

**Figure 4.**
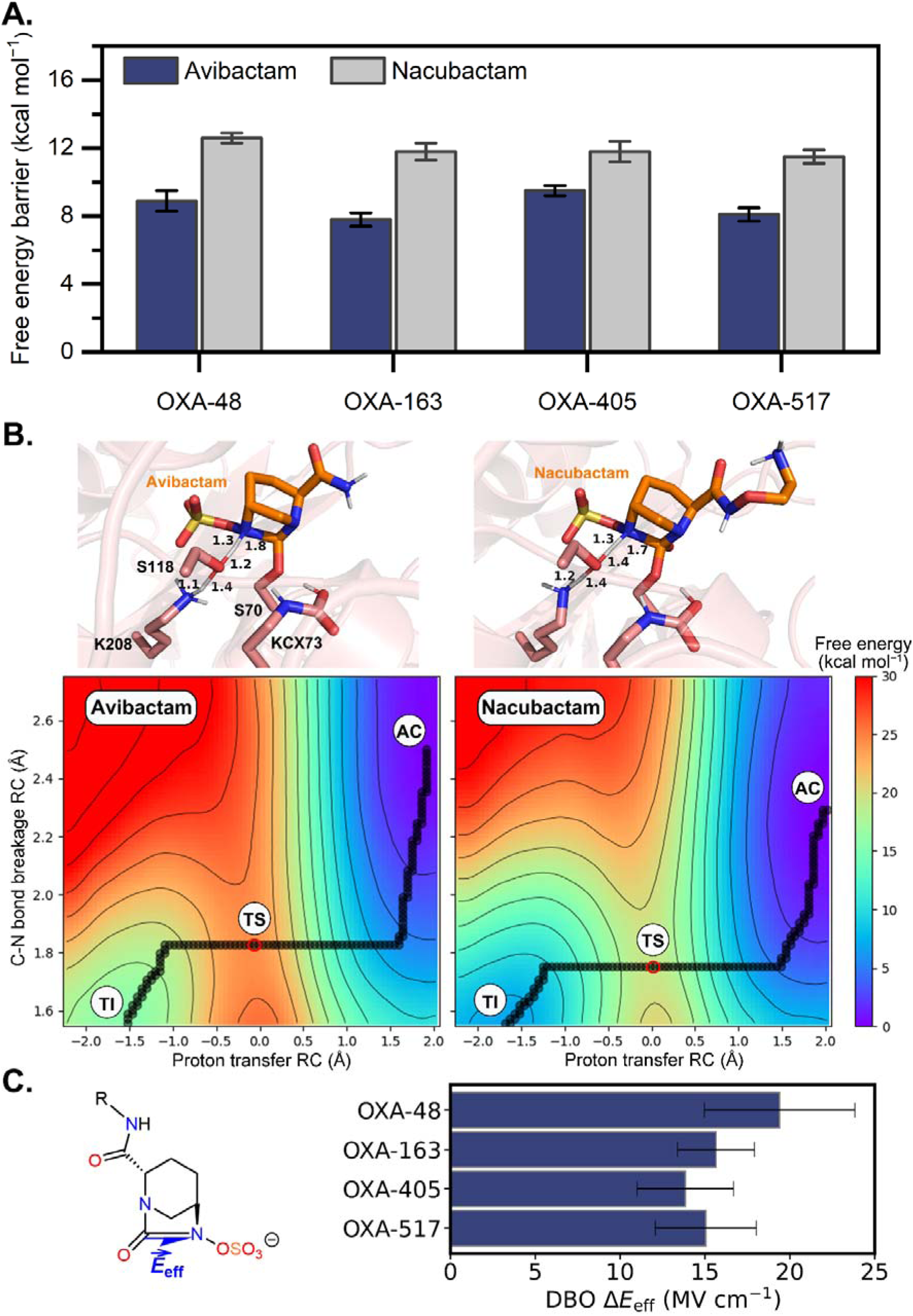
Predicted free energy barriers for the second step of DBO acylation of OXA-48 variants. (A) Free energy barriers at the XTB2/ff14SB level for OXA-48 variants with avibactam and nacubactam. Values are the Boltzmann-weighted barriers from five independent umbrella sampling simulations with error bars indicating the standard deviations. (B) Representative transition state structures and free energy surfaces corresponding to the lowest free energy barrier for the second step of acylation of OXA-48 by avibactam (left panel) and nacubactam (right panel). Minimum free energy paths (MFEPs) indicated by spheres, with AC = acylenzyme, TS = transition state, TI = tetrahedral intermediate. (C) Difference in the effective electric field contributions of nacubactam and avibactam projected along the DBO C-N bond, calculated using the transition state structures and defined as 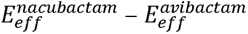. The standard deviation of the difference was calculated using the standard law of propagation of uncertainties 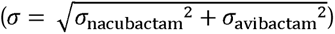.

The transition state for the second acylation step from our QM/MM MD simulations revealed a similar location as in the CI-NEB potential energy profiles. For OXA-48 acylation by avibactam, the transition state was reached when both proton transfers (from and to Ser118) are approximately half-way (proton transfer reaction coordinate approaches 0.0 Å) and the C-N bond distance approaches 1.9 Å (**Figure 4B**, left panel), similar to the CI-NEB results (**Figure 3A**). Equivalent calculations for the reaction of nacubactam with OXA-48 (**Figure 4B**, right panel) showed that the minimum free energy paths and TS geometries are highly similar between both DBOs, and no obvious differences in the active site interactions could be observed. Hence we sought other explanations for the differing behaviours of the two DBOs. Given that the active site electric field has been shown to explain differences in reaction barriers for carbapenem (*β*-lactam) hydrolysis by other serine *β*-lactamases ^52, 64^ and that avibactam and nacubactam have different net charges, we next investigated the possible influence of the electric field from the enzyme-substrate environment on the DBO acylation barrier. The electric field was projected on the DBO C-N bond (**Figure 4C**, left panel), which breaks in the second acylation step, with the nitrogen becoming protonated. The resulting effective electric field was calculated using the sampled transition state ensembles of each enzyme-substrate system. To investigate this more broadly, simulations of the second step of acylation were also performed for the *β_5_*–*β_6_*loop variants OXA-405 and OXA-517. These yielded similar free energy barriers for both DBOs, with the barrier for avibactam again lower (by 2.3 and 3.4 kcal mol^−1^) that for nacubactam for OXA-405 and OXA-517, respectively (**Table S4**).

For all four enzymes, the effective electric field (including solvent) for the reactions with avibactam is more negative than that with nacubactam, consistent with the expected catalytic effect of a (more) negative electric field on the C-N bond breakage. The differences are 15.8, 9.7, 11.8 and 13.6 MV cm^−1^ for OXA-48, -163, -405 and -517 respectively. This means that the magnitude of the difference correlates with the difference in IC_50_ values for OXA-48, - 163 and -405. When the contributions to the effective electric field are decomposed per residue, it is clear that there are no significant differences for the majority of enzyme residues (**Figure S12**). Contributions of the *β_5_*–*β_6_* loop are also similar among all systems (slightly positive), despite the distinct differences in sequence (**Figure 2**). The main difference is that in OXA-163, Asp212 (equivalent to Ser212 in OXA-48) provides a negative contribution to the effective electric field, which is absent in the other variants. However, this does not cause a difference between the transition state ensembles of the second acylation step for avibactam and nacubactam. Instead, the observed difference in electric field between the DBOs in all four enzymes is caused by the DBOs themselves: avibactam consistently has a negative contribution to the effective electric field, whereas that of nacubactam is positive (**Figure 4C**, right panel; **Figure S12**).

The tail groups on the DBO core are highly flexible in the simulations, leading to large fluctuations in the contributions of the DBOs to the effective electric field, but the contribution from nacubactam is consistently positive. A key factor in this contribution is expected to be the terminal NH_3_^+^ moiety. Indeed, when we neutralise this group and repeat the calculations, the difference in effective electric field contributions between avibactam and nacubactam in OXA-48 reduces from 19.4 to 12.4 MV cm^−1^. However, p*K*_a_ predictions based on the MC and AC structure ensembles from our simulations indicate that the terminal group can be assumed to be fully protonated in the active site (p*K*_a_ > 9 in all cases; IC_50_ assays performed at pH 7.6). Thus, the nacubactam tail group, and in particular the NH_3_^+^ moiety, leads to a higher barrier for DBO acylation, thereby resulting in a lower inhibitory effect of nacubactam. Two additional DBO inhibitors (relebactam and zidebactam) that similarly contain ionisable nitrogen groups in their tails also exhibit reduced potency against OXA-48 relative to avibactam ^62, 65^. Together, these observations indicate that ionisable nitrogen groups in the DBO tail generally diminish inhibitory potency against OXA 48 ^30^.

Overall, our reaction simulations and electric field analysis can explain the difference in IC_50_ values for inhibition of the same enzyme by avibactam and nacubactam. However, it is not clear why nacubactam inhibits OXA-48 with a much higher IC_50_ value compared to OXA-163 (and to a lesser extent OXA-405). To address this question further, the non-covalent binding of the two DBO inhibitors with OXA-48-like enzymes is investigated in the following section.

### Changes in the β_5_–β_6_ loop affect nacubactam binding in OXA-48 variants

To investigate whether the non-covalent binding contributes to the distinct differences in inhibition potency between avibactam and nacubactam observed for OXA-48 and -163 (**Figure 2**), 10x 10 ns MM MD simulations of the Michaelis complexes were performed per system, followed by MM-PBSA binding energy calculations on all trajectories. Among the OXA-48-like enzymes, nacubactam consistently exhibits a lower binding affinity (higher binding energy) compared to avibactam (**Figure 5A** and **Table S5**). A significant difference in binding energy was further observed between nacubactam bound to OXA-48 and OXA-163 (**Figure 5A**). Moreover, the difference in binding energy between avibactam and nacubactam is the smallest for OXA-163 (5.2 kcal mol^−1^), and the largest for OXA-48 (7.9 kcal mol^−1^) (**Table S5**). This trend is in agreement with the experimentally observed IC_50_ differences ^30^. To confirm the results from MM-PBSA (an approximate method), relative binding free energy calculations were also performed with the alchemical transfer method ^55, 56^, which showed the same trend between avibactam and nacubactam (**Table S5** and **Figure S13**).

**Figure 5.**
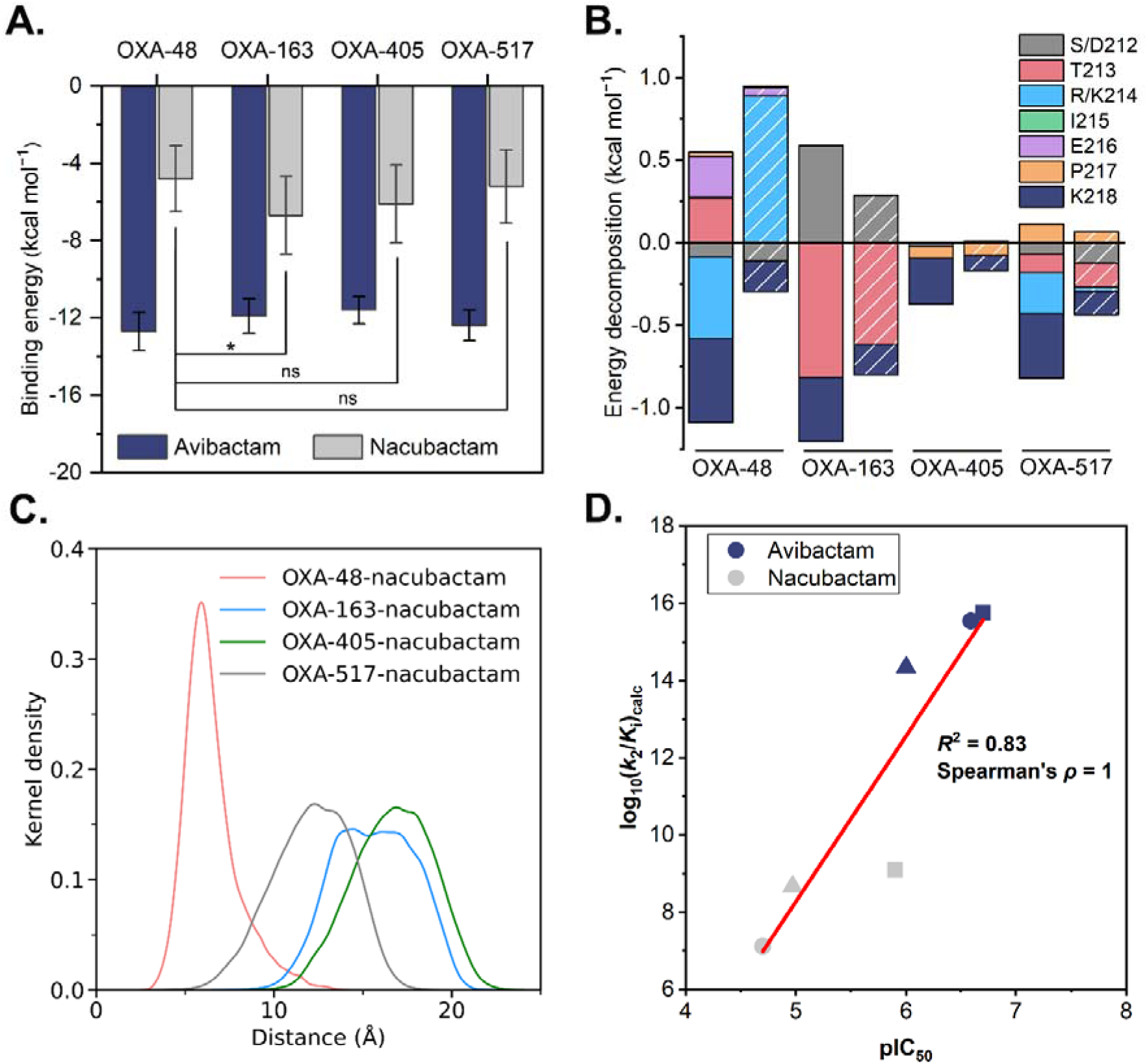
(A) Binding energies for OXA-48 variants with avibactam and nacubactam calculated with MM-PBSA based on 100 ns MM MD simulation of Michaelis complexes (ten replicas of 10 ns per system). The *p* values were obtained from independent two-sample, two-tailed *t*-tests (* *p* < 0.05; ns = not significant). (B) Binding energy decomposition of residues 212 and 213–218 (the *β_5_*–*β_6_*loop). Solid and patterned bars correspond to avibactam- and nacubactam-bound Michaelis complexes, respectively. (C) Distributions (kernel density estimates) of the closest distance between the nacubactam ammonium group and Arg214 in OXA-48, Lys218 in OXA-163 and -405 or Lys214 in OXA-517 (charged nitrogen) during the Michaelis complex MM MD simulations. Detailed distance measurements are shown in Figure S14. (D) Correlation between experimental pIC_50_ (see Figure 2) and calculated log_10_(*k*_2_/*K_i_*_)_ derived from umbrella sampling reaction barriers and MM-PBSA binding energies. Circle: OXA-48; Square: OXA-163; Triangle: OXA-405. The related values are provided in Table S6.

Decomposition of the MM-PBSA binding energies shows that residue Arg214 in OXA-48 contributes differently to the binding of avibactam and nacubactam (**Figure 5B**). For avibactam, the residue contributes to binding (negative energy contribution, i.e. increasing affinity), whereas it does not for nacubactam (strongly positive energy contribution, i.e. decreasing affinity). A similar trend was also observed for Lys214 in OXA-517: Lys214 favours binding of avibactam, whereas its contribution is negligible in the nacubactam-bound complex. Notably, Lys218 consistently contributes to a more negative binding energy with avibactam than with nacubactam across all OXA-48-like enzymes. Other DBO inhibitors with amino-moieties in their tail groups (e.g., relebactam and zidebactam) have also been reported to exhibit lower binding affinities against OXA-48 ^66^, suggesting that protonation of the ionisable tail group could impair binding to OXA-48. Unlike OXA-48, in OXA-163 and - 405, the residue contributions to binding are similar for avibactam and nacubactam. Due to the Ser212Asp substitution and a four-residue deletion in the *β_5_*–*β_6_* loop of OXA-163, Asp212 contributes positively, which is compensated by a negative contribution from Thr213. Thus, the overall contribution of this region favors binding of avibactam and nacubactam to OXA-163.

Inspection of the nacubactam Michaelis complex MD trajectories (**Figure S14**) indicates that for OXA-48 repulsion between Arg214 and the positively charged ammonium group on nacubactam likely cause the large change in the contribution of this residue (compared to avibactam). In contrast, no such electrostatic repulsion exists between Lys218 and the nacubactam ammonium group in OXA-163 and -405. This is clearly reflected in the distances between the positively charged nitrogen atoms in these sidechains and the nacubactam ammonium group, which are predominantly < 10 Å in OXA-48 and > 10 Å in OXA-163 and -405 (**Figure 5C**; trends remain the same in longer MD trajectories, see **Figure S15**). The repulsion between Arg214 and nacubactam is maintained in OXA-48 as Arg214 is anchored in place through a salt-bridge interaction with Asp159. In OXA-517, the equivalent interaction (between Lys214 and Asp159) is occasionally present ^67^, leading to an intermediate distance and thus causing some electrostatic repulsion. Overall, our analysis of non-covalent binding indicates that sequence changes in the *β_5_*–*β_6_* loop can cause significant differences in the strength of interactions of OXA-48 variants with nacubactam, but not avibactam. In particular, the presence of two positively charged residues in this region in OXA-48 and -517 (compared to only Lys218 in OXA-163 and -405) leads to weaker non-covalent binding of nacubactam.

### Comparison with and prediction of experimental data

From the results obtained so far, it is clear that both the barrier for DBO acylation and the initial non-covalent DBO binding can play a role in the effectiveness of inhibition of OXA-48-like *β*-lactamases. Indeed, it is considered that for covalent inhibitors, both processes will likely influence the experimentally measured IC_50_ values ^68^. Experimental data also show that, even when Arg214 is not present, as in OXA-163 and -405, nacubactam remains less potent than avibactam, indicating that the impaired binding caused by Arg214 cannot fully explain the difference in inhibitory potency. In order to compare our calculated values with the IC_50_ values, we therefore first convert the calculated barriers for the rate-limiting second step of acylation (from umbrella sampling calculations) to the rate for covalent bond formation *k*_2_ and the non-covalent binding energies (from MM-PBSA calculations) to the dissociation constant *K*_i_ for the non-covalent Michaelis complexes (**Table S6**). Then, we combine these as log_10_(*k*_2_/*K*_i_) and compare them to the experimental pIC_50_ values. Because the *k*_off_ rates have been reported to be extremely small for acylation of OXA-48 like proteins by DBO ^16, 62, 63, 69^, the relative differences in log_10_(*k*_2_/*K*_i_) values are expected to reflect the differences in inhibitory potency ^68^, allowing direct comparison with experimental pIC_50_ values. Indeed, a clear correlation is obtained (**Figure 5D**). In particular, the calculated order of inhibition efficiency is the same as that observed experimentally (Spearman’s rank = 1). However, the estimated log_10_(*k*_2_/*K*_i_) shows two distinct populations (with the reported Pearson *R*^2^ value therefore not representative of correlation for individual DBOs), reflecting that inhibition by avibactam (compared to nacubactam) is overestimated in our calculations. Furthermore, the correlation is biased (with slope >>1), likely due to MM-PBSA being an approximate method, with its binding energies not directly comparable to *K*_i_ values. However, based on the obtained correlation, IC_50_ values of avibactam and nacubactam can be predicted for OXA-517, a variant for which these values have not yet been determined experimentally. Using the calculated *K*_i_ and *k*_2_ values and the fitted line, IC_50_ values of avibactam and nacubactam for OXA-517 are predicted to be 0.16 ± 0.06 μM and 10.26 ± 0.12 μM, respectively (uncertainties calculated based on propagating the standard deviations between independent replicas of the barriers from US and binding energies from MM-PBSA). This indicates that avibactam is expected to inhibit OXA-517 similarly to OXA-48. However, our predictions are intended primarily to capture the relative inhibitory potency of the two inhibitors toward OXA-517 (or other variants), rather than to provide precise estimates of their experimental IC_50_ values.

For avibactam, all OXA-48-like enzymes share similar IC_50_ values. For nacubactam, the IC_50_ value of OXA-517 is predicted to lie between those of OXA-48 and -163, which is similar to the order of imipenem hydrolysis activities across these enzymes ^29, 67^. Nacubactam exhibits a bulkier molecular structure than avibactam, with larger substrates previously suggested to be less favourably accommodated by OXA-48 than by OXA-163 ^18^. Previous work reported a ceftazidime-carbapenem activity trade-off between OXA-48 and -163, likely due to a combination of increased accessibility of the OXA-163 active site to the bulkier substrate ceftazidime ^24^, as well as subtle changes in active site solvation ^70^. A similar ceftazidime-avibactam trade-off was also observed in OXA-48, where mutations that increase flexibility within the active site enhance hydrolysis of ceftazidime while reducing avibactam acylation activity ^71^. Another study showed that the catalytic efficiency of OXA-163 towards ceftazidime (*k*_cat_/*K*_M_) was substantially higher than the acylation efficiency of avibactam (*k*_2_/*K*) ^69^. Our results show that differences between OXA-48 variants in their susceptibility to DBO inhibitors arise from more than just size-based effects: the positively charged tail group of e.g. nacubactam negatively effects both non-covalent binding and the acylation reaction. We expect similar effects for other DBOs with positively charged tail groups, such as relebactam and zidebactam. Indeed, kinetic data show that relebactam, zidebactam, and a close analogue of zidebactam, all with positive charge in their tail groups, exhibit no measurable inhibitory activity towards OXA-48 ^63^. In contrast, durlobactam, which does not have a positively charged tail group, exhibits substantially stronger inhibition of OXA-48 than nacubactam or zidebactam, and is slightly more effective than avibactam ^62^. Similarly, another novel DBO inhibitor without positive charge, WCK 4234, also exhibits greater inhibitory potency towards OXA-48 than relebactam and zidebactam, according to kinetic data ^63^. Based on our findings, the expectation is that such DBOs without positive charged tail groups will be similarly effective as inhibitors in all OXA-48 variants studied here.

## Conclusion

Due to the growing threat of antimicrobial resistance, there is an urgent need to develop antibacterials with novel mechanisms of action, including combinations of *β*-lactams and new types of *β*-lactamase inhibitors ^5, 72^. DBOs, a new class of non-*β*-lactam *β*-lactamase inhibitors, are attracting increasing attention. Understanding the inhibition mechanism of DBO inhibitors, and the basis for their differential inhibitory effects towards different *β*-lactamase targets, should help combat *β*-lactamase-mediated antibiotic resistance. In this work, we reveal the detailed mechanism of DBO acylation across different OXA-48-like enzymes, including cephalosporin-hydrolysing variants, and with different DBOs ^22^. Despite differences in the active site structural environment and enzyme-inhibitor interactions, DBO acylation was found to proceed through the same chemical mechanism across different enzyme-inhibitor pairs. Moreover, the second step of DBO acylation was established as rate-determining, thereby causing differences in acylation efficiency between avibactam and nacubactam. This difference is related to differences in the active site electric field, with the positive charge on the nacubactam tail negatively influencing acylation efficiency. Based on available data of inhibition with other DBOs ^62, 63^, this effect of charged tails is expected to be a major influence on acylation efficiency in OXA-48-like enzymes.

Non-covalent binding further affects the inhibition efficiency (and thus IC_50_) of DBOs. Nacubactam was predicted to have significantly lower binding affinity than avibactam, particularly for OXA-48, consistent with the measured difference in IC_50_ for these DBOs ^30^. This reduced binding affinity is attributed to electrostatic repulsion between positively charged residues in the active site and nacubactam, with Arg214 in OXA-48 having a particularly strong effect. The absence or substitution of this residue in other OXA-48-like variants reduces the unfavourable interactions with nacubactam.

By combining the calculated non-covalent binding affinities and reaction barriers for acylation (covalent binding), a good correlation with experimental IC_50_ values was obtained, with the overall experimental trend being reproduced. This correlation suggests that our simulations can be used to predict the (relative) DBO inhibition efficiency for other OXA-48-like enzymes, as we demonstrate with OXA-517. Our work thereby highlights the potential of computational approaches for evaluation of covalent inhibitors by accounting for both their non-covalent interactions and covalent reactivity. Previously, similar approaches have been applied to covalent inhibitors targeting epidermal growth factor receptor (EGFR) ^73^ and Bruton tyrosine kinase (BTK) ^74^, for example.

In summary, we used multiscale simulations to reveal both the acylation reaction and non-covalent binding between DBOs and OXA-48-like enzymes in atomic detail. We establish that both non-covalent binding and reaction barriers contribute to the experimentally measured differences in inhibitory potency between avibactam and nacubactam. Our results thus provide an in-depth understanding of DBO inhibition of OXA-48-like enzymes, and explain the substantial differences observed in experimental inhibitory effects, highlighting the roles of active site electric field and electrostatics in modulating inhibitor potency. Overall, this work shows how multiscale simulations can be used to determine to the detailed molecular origins of serine *β*-lactamase inhibition with covalent inhibitors such as DBOs. Therefore, our simulation protocols can help to anticipate DBO effectiveness against new *β*-lactamase variants, and inform further DBO development.

## Data availability

The input files, including the settings and initial structures for all simulations and QM/MM potential energy calculations, as well as the parameters for the carboxylated Lys73, and avibactam and nacubactam in the free-ligand form, will be made freely available online upon publication.

## Author contributions

DW and MWvdK conceived the work. DW performed all simulations, with assistance from RC, and initial work performed by VHAH. DW and MWvdK performed analysis and interpretation of results. MWvdK supervised DW, RC and VHAH, with contributions from JS. DW and MWvdK wrote the manuscript draft, with revisions from JS and approval by all authors.

## Notes

The authors declare no competing financial interest.

## Acknowledgments

We thank the GW4 BIOMED2 DTP for DW’s studentship, grant MR/W006308/1, and the GW4 BIOMED DTP for VHAH’s studentship, grant MR/N0137941/1, awarded to the Universities of Bath, Bristol, Cardiff and Exeter from the Medical Research Council (MRC)/UKRI. JS acknowledges funding from the U.K. Biotechnology and Biological Sciences Research Council (BBSRC, BB/W001187/1) and the European Research Council under the European Horizon 2020 research and innovation program (PREDACTED Advanced Grant Agreement no. 101021207). All simulations and QM/MM potential energy calculations were conducted using the facilities of the Advanced Computing Research Centre at the University of Bristol (https://www.bris.ac.uk/acrc/).

