## Supporting Information for "Active site electrostatic effects modulate DBO inhibitor potency against OXA-48 variants"

This Supporting Information contains:

- Detailed description of tested semi-empirical QM methods for umbrella sampling, applied restraints for simulations and settings for relative binding free energy calculations (Notes S1–S4);
- Detailed definition of the QM region used in the reaction simulations (Figure S1);
- Visualization and quantification of different active-site hydration states during Michaelis complex MM MD simulations (Figure S2, Table S1);
- Single-point energies of the QM regions calculated using different semi-empirical QM methods compared with high-level DFT (Figures S3–S4);
- Detailed minimum energy pathways and potential energy profiles of acylation of OXA-48 or OXA-163 by avibactam with different active-site hydration states and QM region definitions (Figures S5–S10);
- Representative transition state structures and free energy surfaces for the first step of acylation (Figure S11);
- Detailed per residue contributions to the effective electric field at the transition states for the second step of acylation (Figure S12);

- Visualization of different equilibrated nacubactam-bound OXA-48 Michaelis complex structures from relative binding free energy and MM-PBSA calculations (Figure S13);
- Electrostatic repulsion of Michaelis complex MM MD simulations (Figures S14–S15);
- Calculated free energy barriers of acylation of OXA-48-like enzymes by DBOs (Tables S2–S4);
- Calculated binding energies from MM-PBSA and relative binding free energies for DBOs non-covalently bound to OXA-48-like enzymes (Table S5);
- Calculated $\log_{10}(k_2/K_i)$ derived from umbrella sampling reaction barriers and MM-PBSA binding energies for the acylation of OXA-48-like enzymes by DBOs (Table S6).

**Note S1**

In order to save the computational cost and identify the best semi-empirical method for QM/MM MD simulation, the QM regions of the images from both steps of CI-NEB calculation of OXA-48 with avibactam were extracted, and single point energy calculations were performed using SCC-DFTB (DFTB2) [1], DFTB3 [2], RM1 [3], PDDG-PM3 [4], PM6 [5] in the sqm program [6] of AmberTools24 [7]. The obtained potential energy profiles were compared to those at the r2SCAN-3c [8] and ωB97X/def2-TZVP [9, 10] levels from ORCA 6.0 [11] to determine the best semi-empirical method for the rate-limiting step simulation. The GFN2-xTB (XTB2) [12] method, as implemented in ORCA 6.0, was also tested for comparison.

**Note S2**

In order to preserve the reactive conformation of Michaelis complex, several additional restraints were applied during the 50 ps QM/MM MD relaxation. Those additional restraints include a one-sided harmonic distance restraint to keep the distance between the Ser118 hydroxyl hydrogen and the DBO $\gamma$-lactam ring nitrogen involved in the C-N bond cleavage below 2.5 Å; a one-sided harmonic distance restraint to maintain the Ser118 hydroxyl O-H bond length below 1.0 Å; three one-sided harmonic distance restraints between the oxygens of the DBO sulfate ester group and the Ser118 hydroxyl hydrogen to prevent the distance from dropping below 2.5 Å; a one-sided harmonic distance restraint to keep the distance between the Ser70 hydroxyl oxygen and the electrophilic carbon above 2.0 Å; two one-sided dihedral angle restraints to ensure that the DBO $\gamma$-lactam ring nitrogen points toward Ser118 and that the DBO sulfate ester group adopts a conformation close to the crystal structure; a one-sided dihedral angle restraint on Lys208 $\chi_3$ to keep the dihedral angle below 100°; and a two-sided harmonic dihedral angle restraint on the diazabicyclooctane ring to maintain its chair conformation. The force constant for all restraints was set to 100 kcal mol$^{-1}$ Å$^{-2}$ or 100 kcal mol$^{-1}$ rad$^{-2}$.

**Note S3**

Two reaction coordinates were applied during QM/MM US MD simulation of the first step of acylation of OXA-48 and OXA-163 by DBOs. One reaction coordinate corresponds to the proton transfer coordinate between KCX and Ser70, defined as distance ($d$[KCX@OQ1, Ser70@H] − $d$[Ser70@O, Ser70@H]) which shows the break of O-H bond in Ser70 and formation of KCX@OQ1-Ser70@H bond. The other is the nucleophilic attack of the $OH^-$ of Ser70 on the electrophilic carbon of DBO, defined as (*d*[Ser70@O, DBO@C]). The reaction was simulated in the reverse direction, from the tetrahedral intermediate to the Michaelis complex. Hence, the proton transfer coordinate value changes from 0.6 Å to −0.6 Å, while for nucleophilic attack, it changes from 1.5 Å to 3.0 Å (both in steps of 0.1 Å). Similar restraints to those used in the second step QM/MM US MD simulation were applied: only the two-sided dihedral angle restraint used to maintain the DBO in chair conformation, those one-sided distance restraints used to prevent proton transfer from Lys208 to the DBO sulfate ester group and a one-sided distance restraint applied to the Ser118 hydroxyl O-H to maintain the distance below 1.0 Å. 208 US windows were used in total for each replica.

**Note S4**

For relative binding free energy calculation, after obtaining the initial structures from MM MD simulations, any water molecules that are closer than 20 Å to the nacubactam carbonyl oxygen were kept. Avibactam was translated by 30 Å so as to be positioned outside of the binding site. The whole system was then solvated with a rectangular box of TIP3P water, with a length of 75 Å in each dimension and was neutralized with $Na^+$ ions. The bonded parameters, atom types and partial charges were retained. The following procedures were then performed: minimisation (10000 steps), equilibration (100 ps), annealing (100 ps, introducing the ATM force), post-annealing equilibration (100 ps, $\lambda$ = 0.5), production (5 ns per $\lambda$ window, 22 $\lambda$ windows). To maintain a conformation similar to that in MM MD simulations, positional flat-bottom restraints were applied to protein Cα atoms, Ser70 and Tyr211 backbone amide groups, the KCX carboxylate group and Ser70 sidechain heavy atoms, and DBO $\gamma$-lactam ring in all procedures. The width of the flat-bottom restraints was set to 0.2 Å, using a force constant of 50 kcal $mol^{-1}$ $Å^{-2}$. A timestep of 2 fs was used. To soften entropic bottlenecks along the alchemical thermodynamic path, the same values of $\alpha$ and uh as in the tutorial were used during production ($\alpha$ = 0.10 kcal $mol^{-1}$ and uh = 110 kcal $mol^{-1}$). The remaining parameters, including the soft-score potential, the protein-ligand centre of mass restraint and the ligand alignment restraint were set to their default values. The perturbation energy values were saved every 100 steps (resulting in 25000 data points per $\lambda$ window), and analysed by using the Unbinned Weighted Histogram Analysis Method (UWHAM) [13] after discarding first 2 ns of the trajectory (15000 data points per $\lambda$ window were used for analysis).

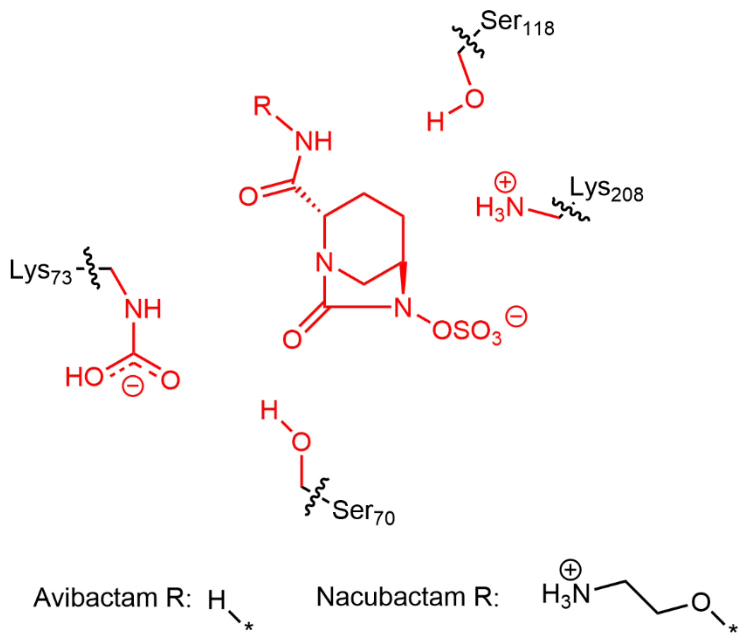


**Figure S1.** QM region in QM/MM MD reaction simulations and potential energy calculations. The whole structure of DBO is included in the QM region. The atoms of QM region are coloured in red, and the breaks between QM and MM region are shown with wavy lines.

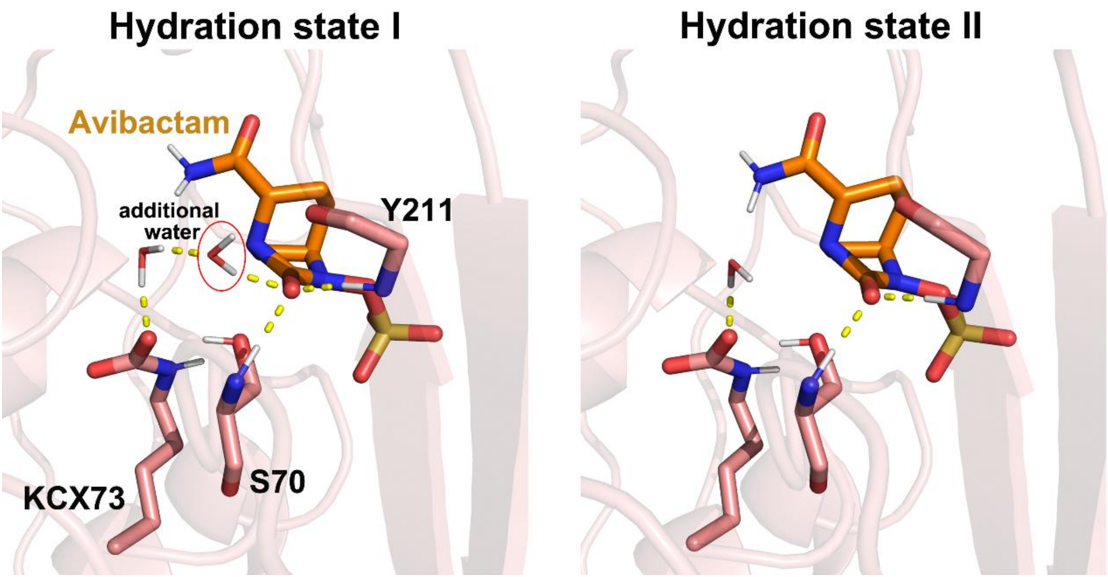


**Figure S2.** Two different hydration states around the avibactam moiety observed during the MM MD and QM/MM MD simulations of the Michaelis complex. The sidechains of carboxylated Lys73 (KCX) (pink), whole Ser70 (pink), the backbone of Tyr211 (pink), active site water molecules and avibactam (orange) were shown as sticks. H-bonds are shown as yellow dashed lines. Snapshots from the 100 ps QM/MM MD of avibactam-OXA-48 were selected for visualization.

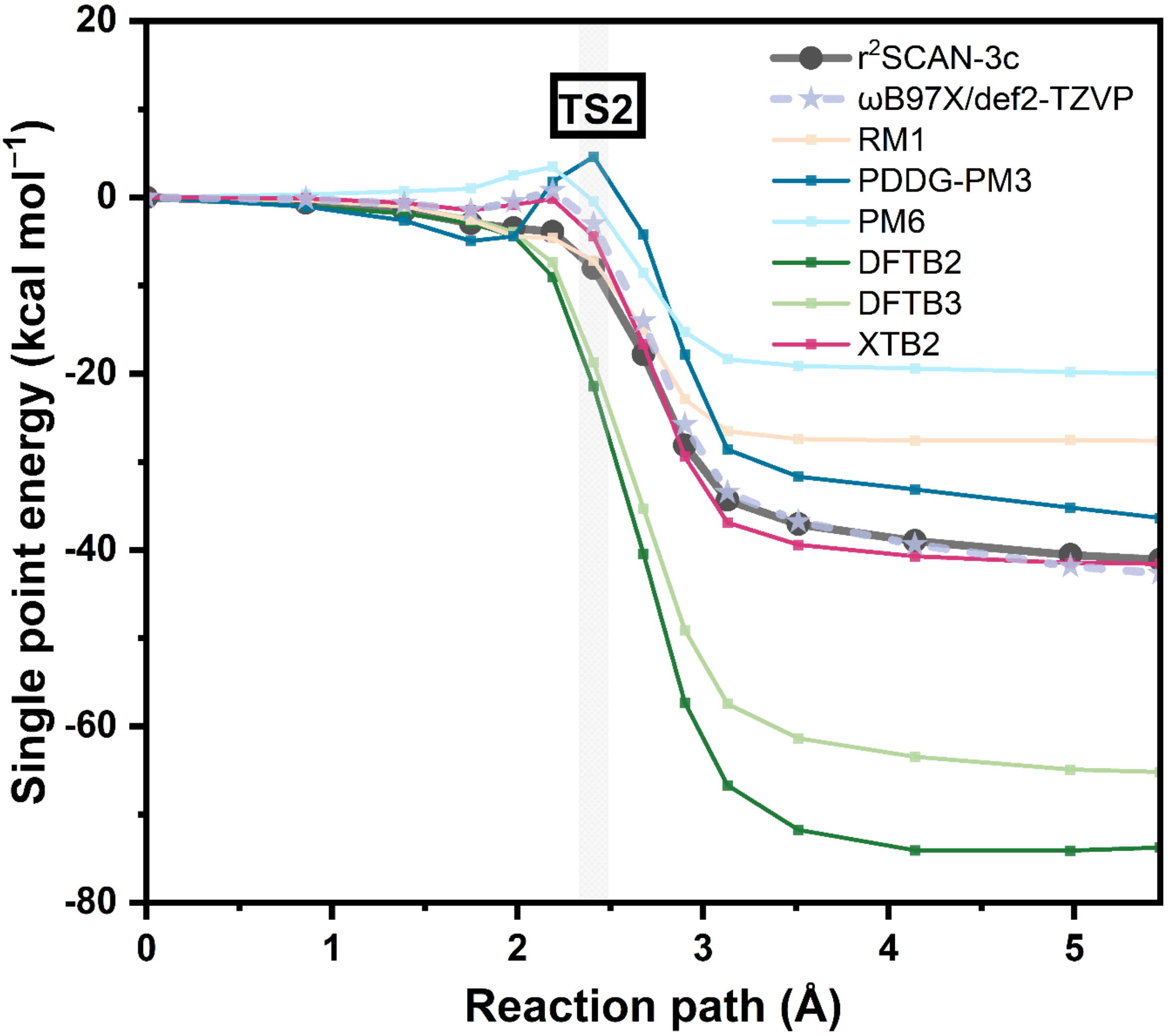


**Figure S3.** Single point energies of the QM regions (with link atoms) of the images from the CI-NEB calculation corresponding to the second step (tetrahedral intermediate → acylenzyme) of avibactam acylation of OXA-48. Energies are reported for different Hamiltonians.

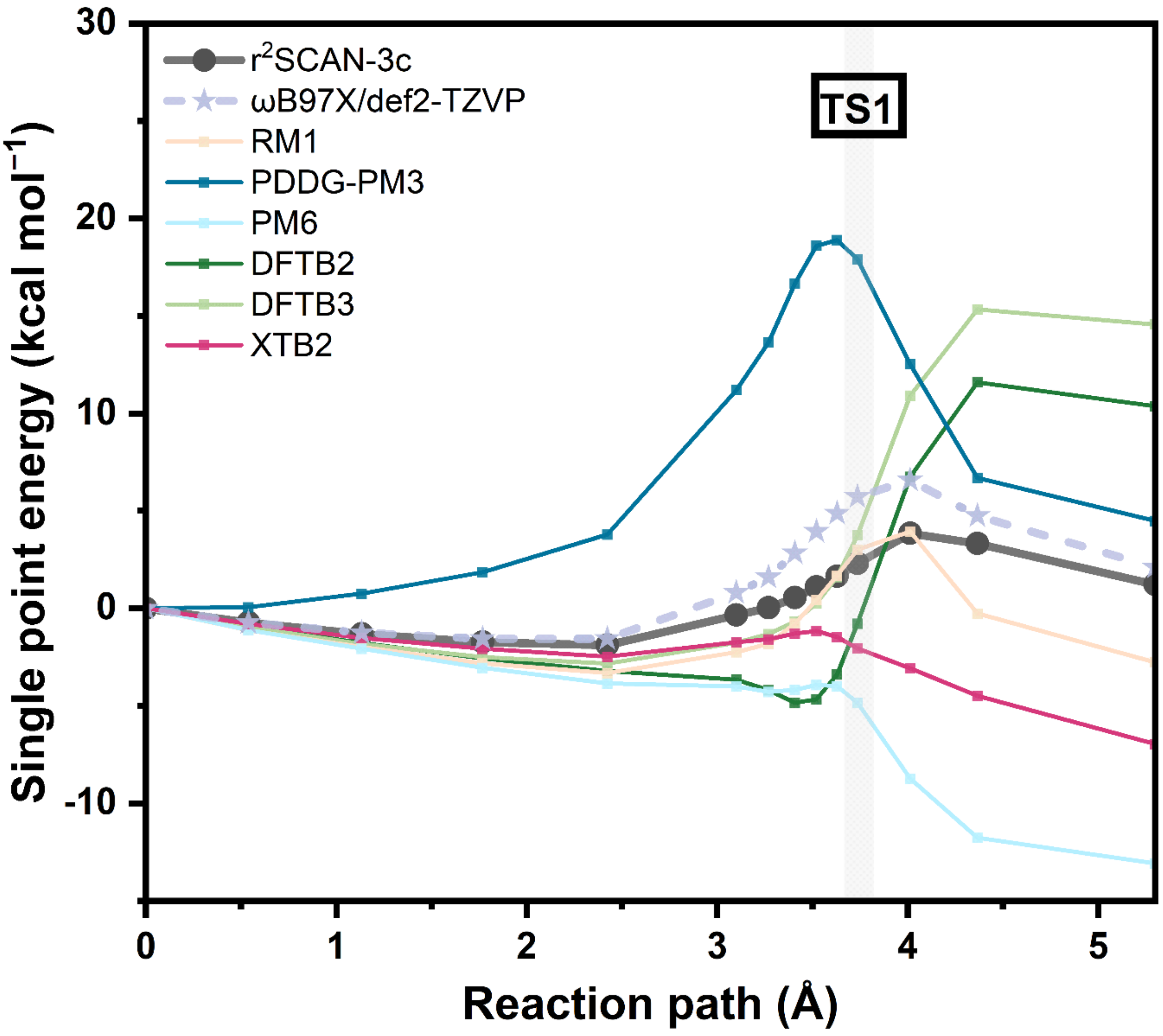


**Figure S4.** Single point energies of the QM regions (with link atoms) of the images from the CI-NEB calculation corresponding to the first step (Michaelis complex → tetrahedral intermediate) of avibactam acylation of OXA-48. Energies are reported for different Hamiltonians.

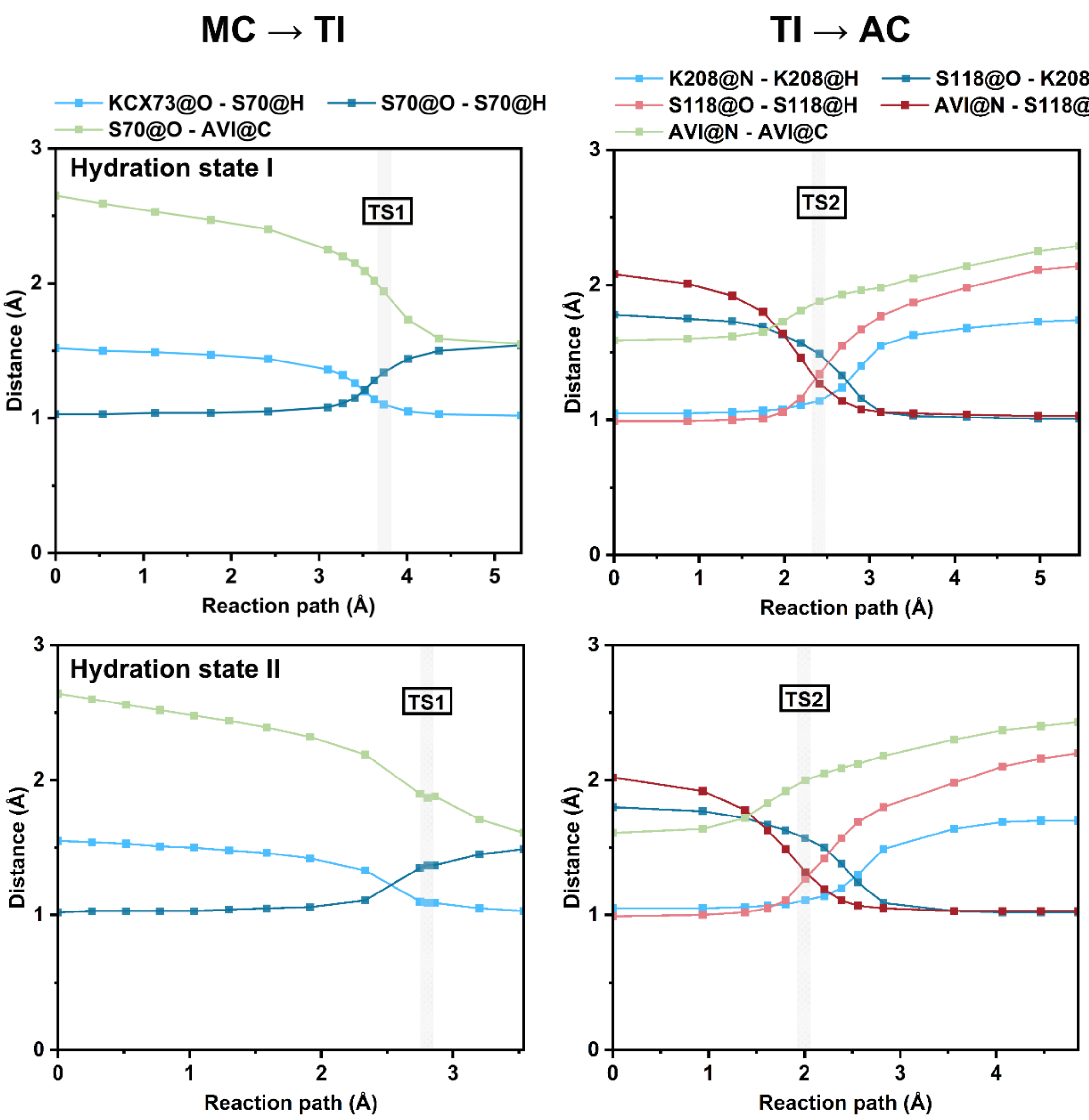


**Figure S5.** Distances of changing bonds on the minimum energy path of avibactam acylation of OXA-48 for TI formation (from MC to TI, left panel), and subsequent AC formation (from TI to AC, right panel) in hydration states I (top panel) and II (bottom panel). MC = Michaelis complex, AC = acylenzyme, TI = tetrahedral intermediate, TS = transition state. CI-NEB calculations were performed at the $r^2$SCAN-3c/ff14SB level.

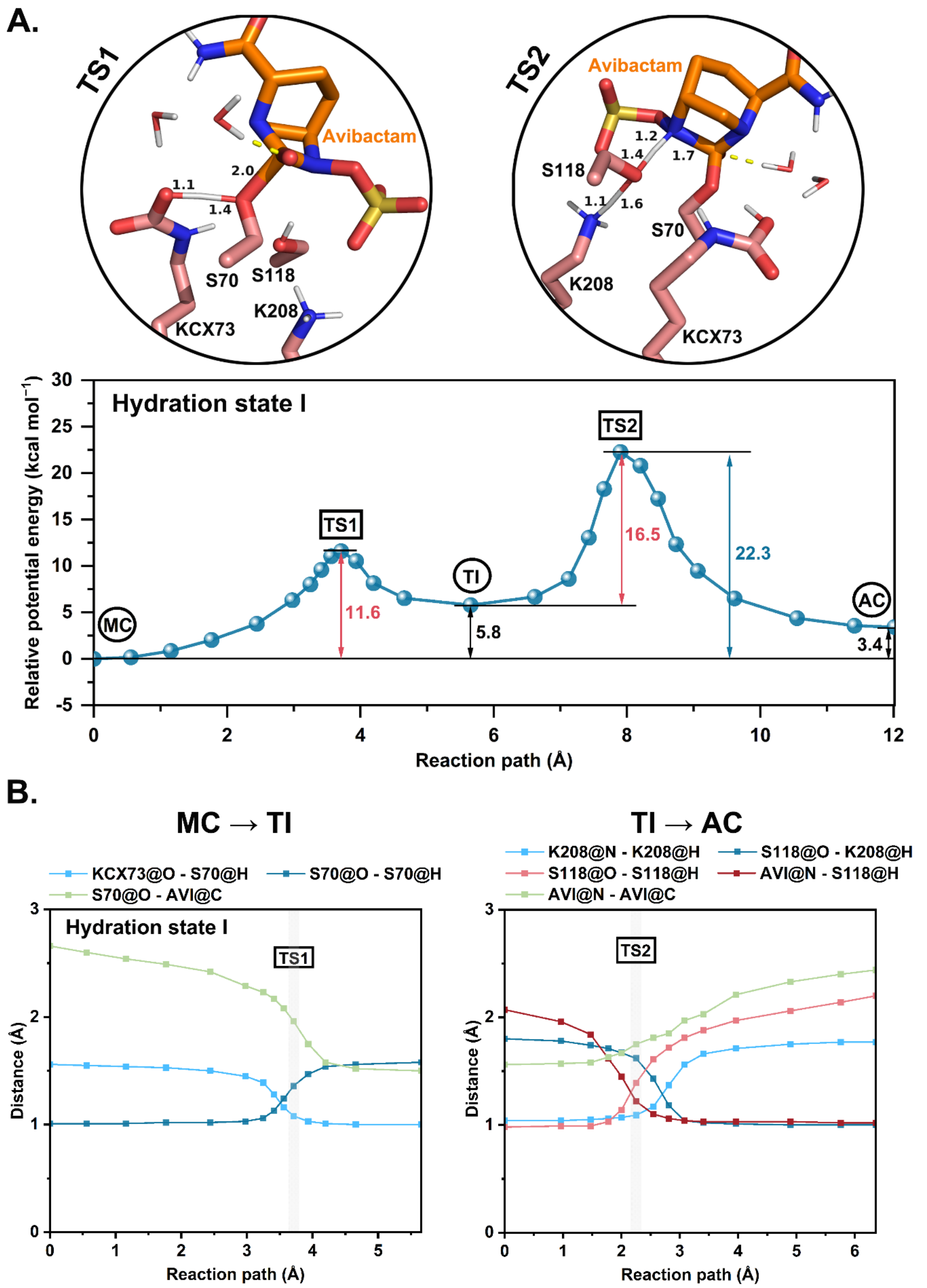


**Figure S6.** Potential energy profile of avibactam acylation of OXA-48 along the minimum energy path in hydration state I, with MC = Michaelis complex, AC = acylenzyme, TI = tetrahedral intermediate, TS = transition state. (A) Potential energy profile of avibactam acylation of OXA-48. CI-NEB calculations, as well as the potential energies, were reported at the ωB97X/def2-TZVP//ff14SB level. The relative energies were determined using MC as the reference. The potential energy barriers for both steps are shown in red, and the overall barrier in blue, while the energies of TI and AC relative to MC are shown in black. The structures of transition states (climbing images) are shown, with reactive residue sidechains (pink carbons) and avibactam (orange carbons) represented as sticks. Distances of changing bonds are labelled in Å. The solvent H-bond to avibactam carbonyl oxygen is indicated by a yellow dashed line. (B) Distances of changing bonds along the minimum energy path shown in (A) for the TI formation (from MC to TI, left panel), and the subsequent AC formation (from TI to AC, right panel) in hydration state I.

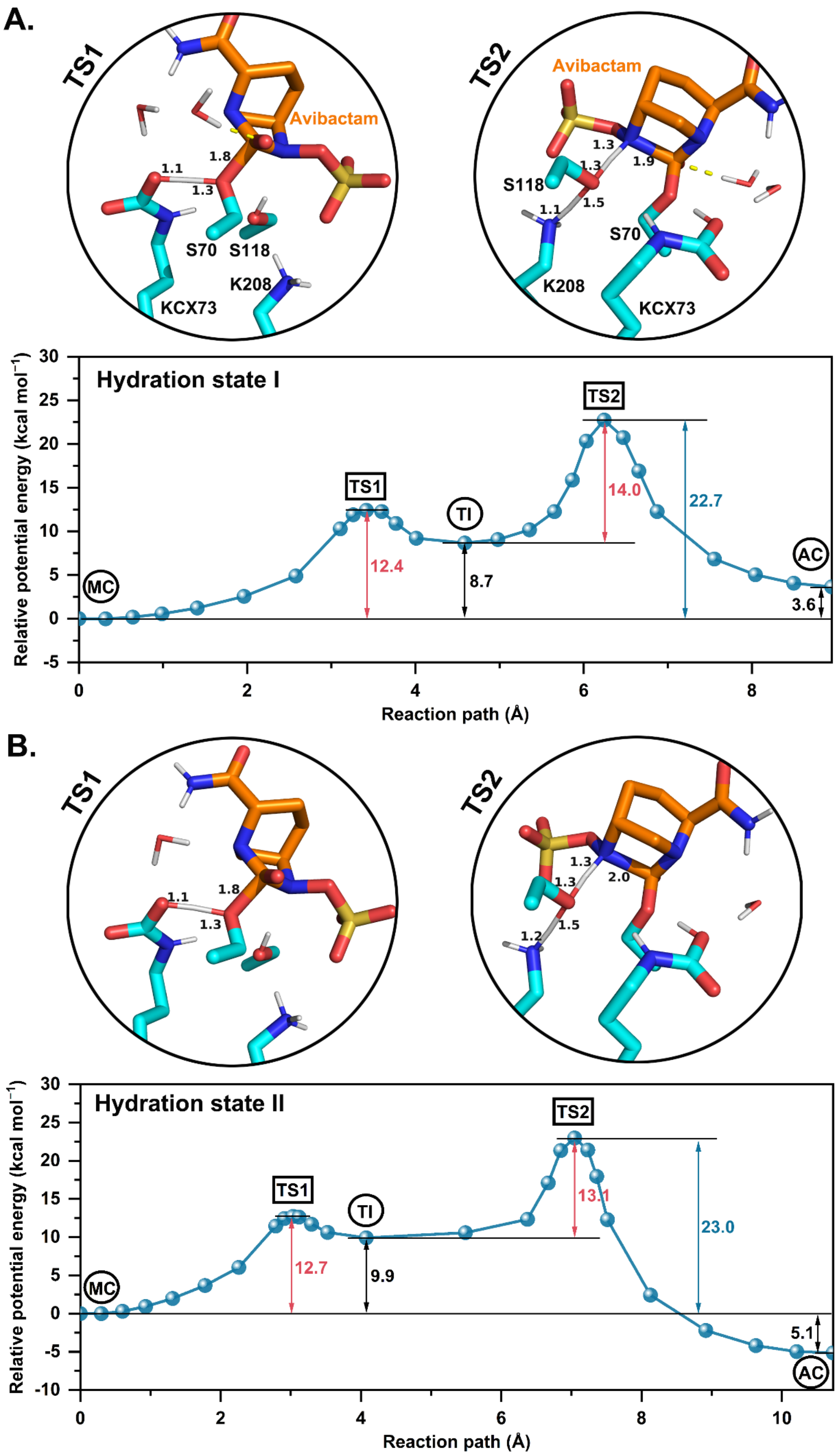
A.
TS1
Avibactam
1.8
1.1
1.3
S70
S118
K208
KCX73
TS2
Avibactam
1.3
1.3
1.9
S118
1.1
1.5
S70
K208
KCX73
Relative potential energy (kcal mol−1)
Hydration state I
TS2
TS1
TI
MC
AC
12.4
8.7
14.0
22.7
3.6
Reaction path (Å)
B.
TS1
1.1
1.8
1.3
TS2
1.3
2.0
1.3
1.2
1.5
Relative potential energy (kcal mol−1)
Hydration state II
TS2
TS1
TI
MC
AC
12.7
9.9
13.1
23.0
5.1
Reaction path (Å)

**Figure S7.** Potential energy profiles of avibactam acylation of OXA-163 in hydration states I (A) and II (B), with MC = Michaelis complex, AC = acylenzyme, TI = tetrahedral intermediate, TS = transition state. CI-NEB calculations were performed at the $r^2$SCAN-3c/ff14SB level, with potential energies refined to the ωB97X/def2-TZVP//ff14SB level through single point energy calculations. The relative energies were determined using MC as the reference. The potential energy barriers for both steps are shown in red, and the overall barrier in blue, while the energies of TI and AC relative to MC are shown in black. The structures of the transition states (climbing images) are shown, with reactive residue sidechains (cyan carbons) and avibactam (orange carbons) represented as sticks. Distances of changing bonds are labelled in Å. The solvent H-bond to avibactam carbonyl oxygen is indicated by a yellow dashed line.

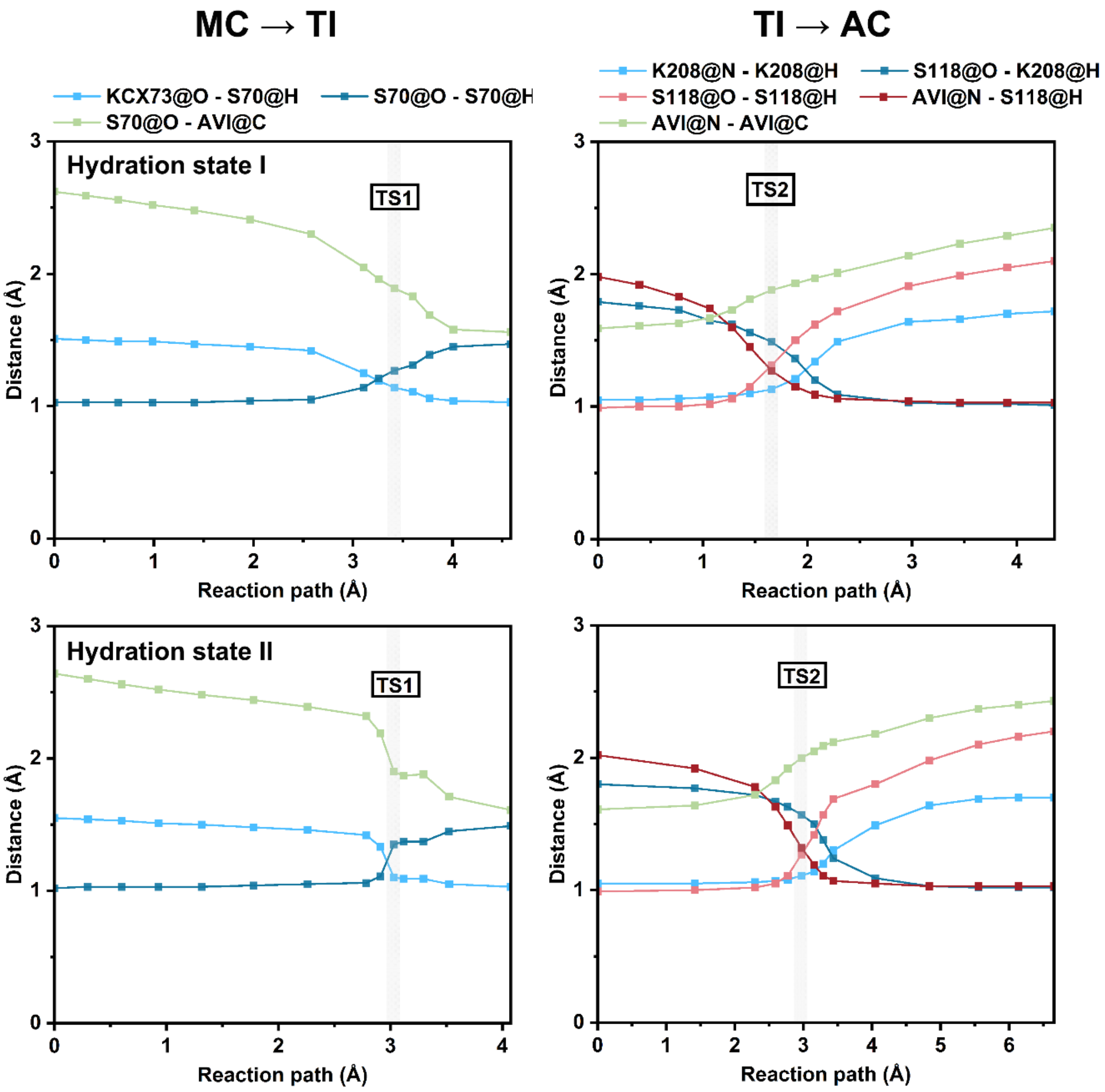


**Figure S8.** The minimum energy path of avibactam acylation of OXA-163 for the TI formation (from MC to TI, left panel), and the subsequent AC formation (from TI to AC, right panel) in hydration states I (top panel) and II (bottom panel), with MC = Michaelis complex, AC = acylenzyme, TI = tetrahedral intermediate, TS = transition state. CI-NEB calculations were performed at the r$^2$SCAN-3c/ff14SB level. Distances of changing bonds in both steps are plotted.

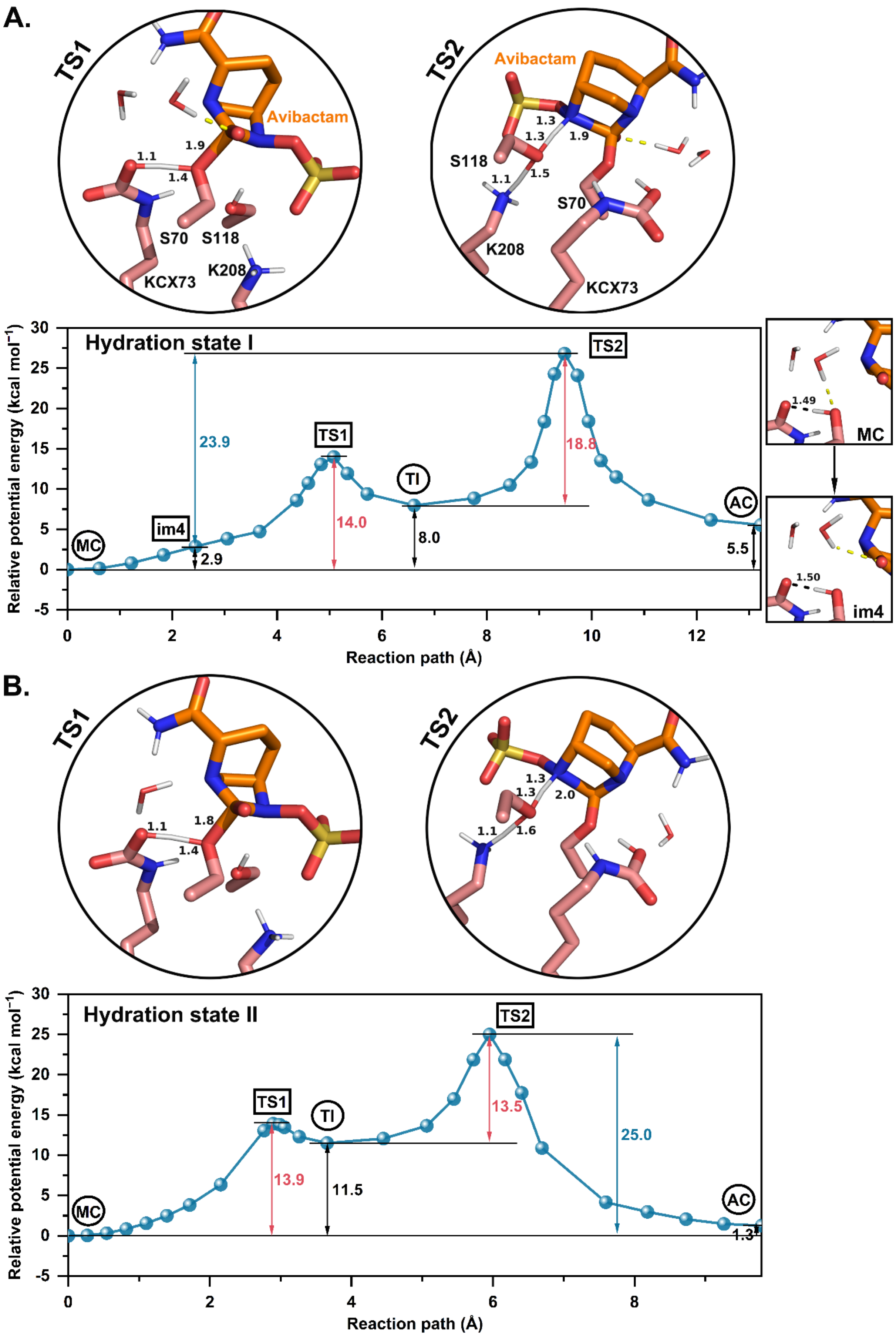

A.
TS1
Avibactam
1.9
1.1
1.4
S70
S118
KCX73
K208
TS2
Avibactam
1.3
1.3
1.9
S118
1.1
1.5
S70
K208
KCX73
Hydration state I
Relative potential energy (kcal mol−1)
Reaction path (Å)
MC
im4
TS1
TI
TS2
AC
23.9
14.0
18.8
8.0
2.9
5.5
1.49
1.50
B.
TS1
1.1
1.8
1.4
TS2
1.3
1.3
2.0
1.1
1.6
Hydration state II
Relative potential energy (kcal mol−1)
Reaction path (Å)
MC
TS1
TI
TS2
AC
13.9
11.5
13.5
25.0
1.3

**Figure S9.** Potential energy profiles of avibactam acylation of OXA-48 in hydration states I (A) and II (B), with MC = Michaelis complex, AC = acylenzyme, TI = tetrahedral intermediate, TS = transition state. Water molecules forming H-bonds with the avibactam carbonyl oxygen and the reactive oxygen of carboxylated Lys73 were included in the QM region. CI-NEB calculations were performed at the $r^2$SCAN-3c/ff14SB level, with potential energies refined to the ωB97X/def2-TZVP//ff14SB level through single point energy calculations. The relative energies were determined using MC as the reference. The potential energy barriers for both steps are shown in red, and the overall barrier in blue (between im4 and TS2 in hydration state I), while the energies of TI and AC relative to MC are shown in black. The structures of the transition states (climbing images) are shown, with reactive residue sidechains (pink carbons) and avibactam (orange carbons) represented as sticks. Distances of changing bonds are labelled in Å. The solvent H-bond to avibactam carbonyl oxygen is indicated by a yellow dashed line.

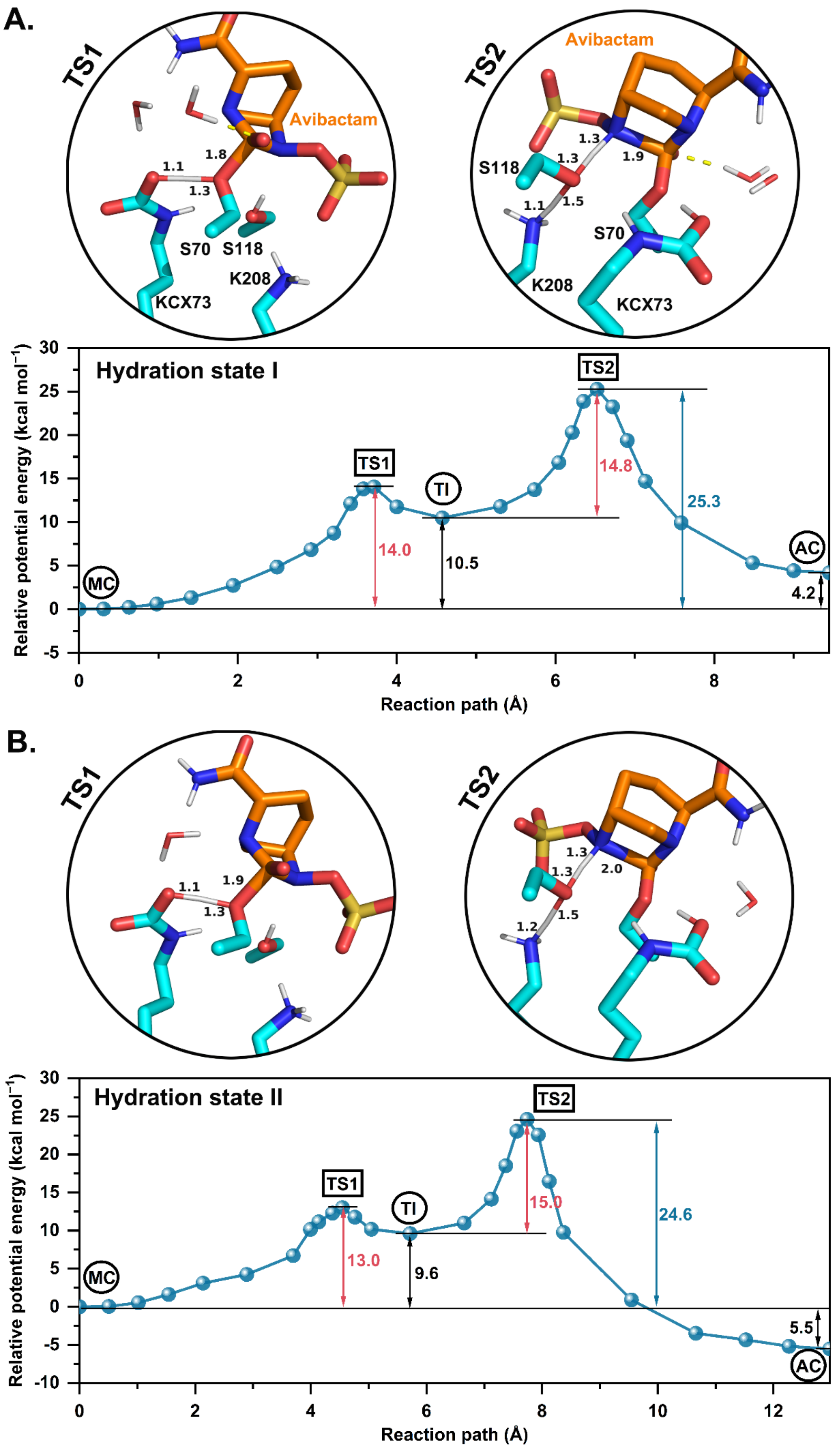
A.
TS1
Avibactam
1.8
1.1
1.3
S70
S118
K208
KCX73
TS2
Avibactam
1.3
1.3
1.9
S118
1.1
1.5
S70
K208
KCX73
Hydration state I
TS2
TS1
TI
14.8
25.3
14.0
10.5
MC
AC
4.2
Relative potential energy (kcal mol−1)
Reaction path (Å)
B.
TS1
1.1
1.9
1.3
TS2
1.3
1.3
2.0
1.2
1.5
Hydration state II
TS2
TS1
TI
15.0
24.6
13.0
9.6
MC
AC
5.5
Relative potential energy (kcal mol−1)
Reaction path (Å)

**Figure S10.** Potential energy profiles of avibactam acylation of OXA-163 in hydration states I (A) and II (B), with MC = Michaelis complex, AC = acylenzyme, TI = tetrahedral intermediate, TS = transition state. Water molecules forming H-bonds with the avibactam carbonyl oxygen and the reactive oxygen of carboxylated Lys73 were included in the QM region. CI-NEB calculations were performed at the r$^2$SCAN-3c/ff14SB level, with potential energies refined to the ωB97X/def2-TZVP//ff14SB level through single point energy calculations. The relative energies were determined using MC as the reference. The potential energy barriers for both steps are shown in red, and the overall barrier in blue, while the energies of TI and AC relative to MC are shown in black. The structures of the transition states (climbing images) are shown, with reactive residue sidechains (cyan carbons) and avibactam (orange carbons) represented as sticks. Distances of changing bonds are labelled in Å. The solvent H-bond to avibactam carbonyl oxygen is indicated by a yellow dashed line.

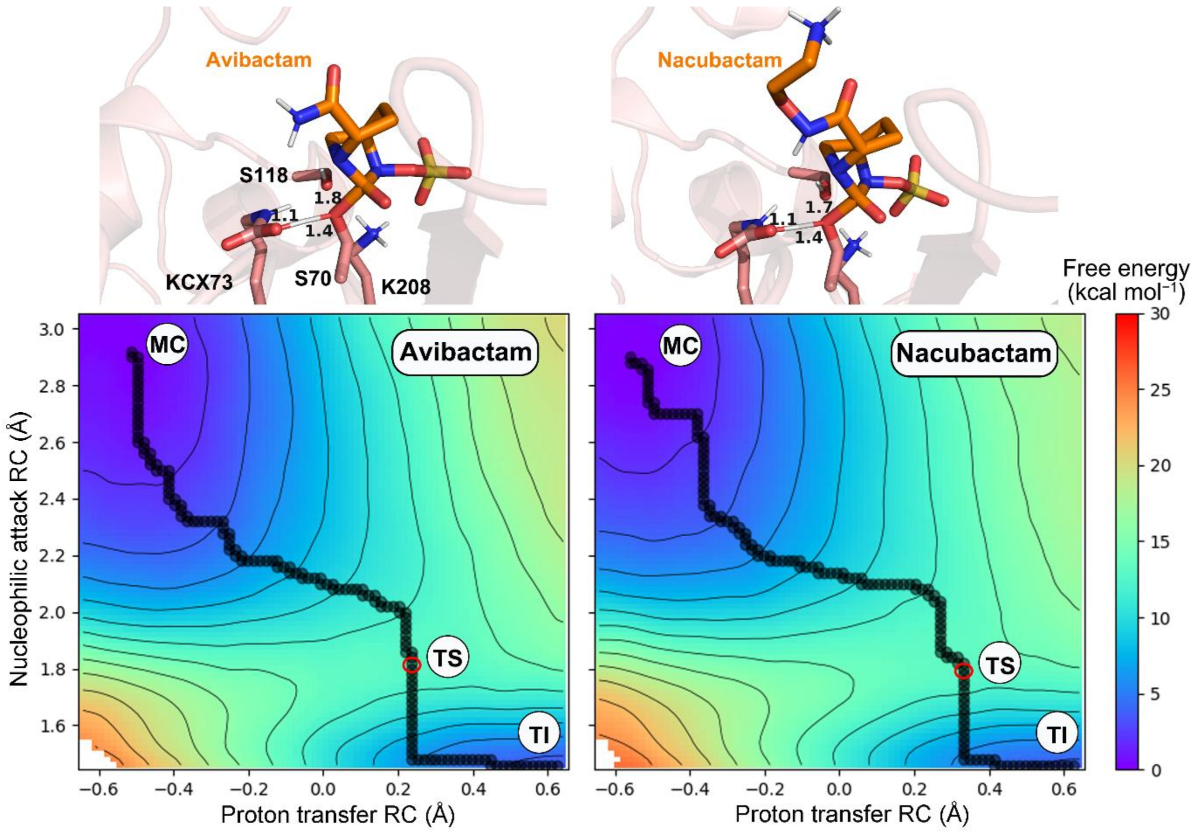


**Figure S11.** Representative transition state structures and free energy surfaces corresponding to the replica with the lowest free energy barrier for the first step of acylation of OXA-48 by avibactam (left panel) and nacubactam (right panel). Obtained through 2D QM/MM umbrella sampling at the RM1/ff14SB level (see Note S3 for details). Minimum free energy paths (MFEPs) are indicated by overlapping black spheres, with MC = Michaelis complex, TS = transition state, TI = tetrahedral intermediate.

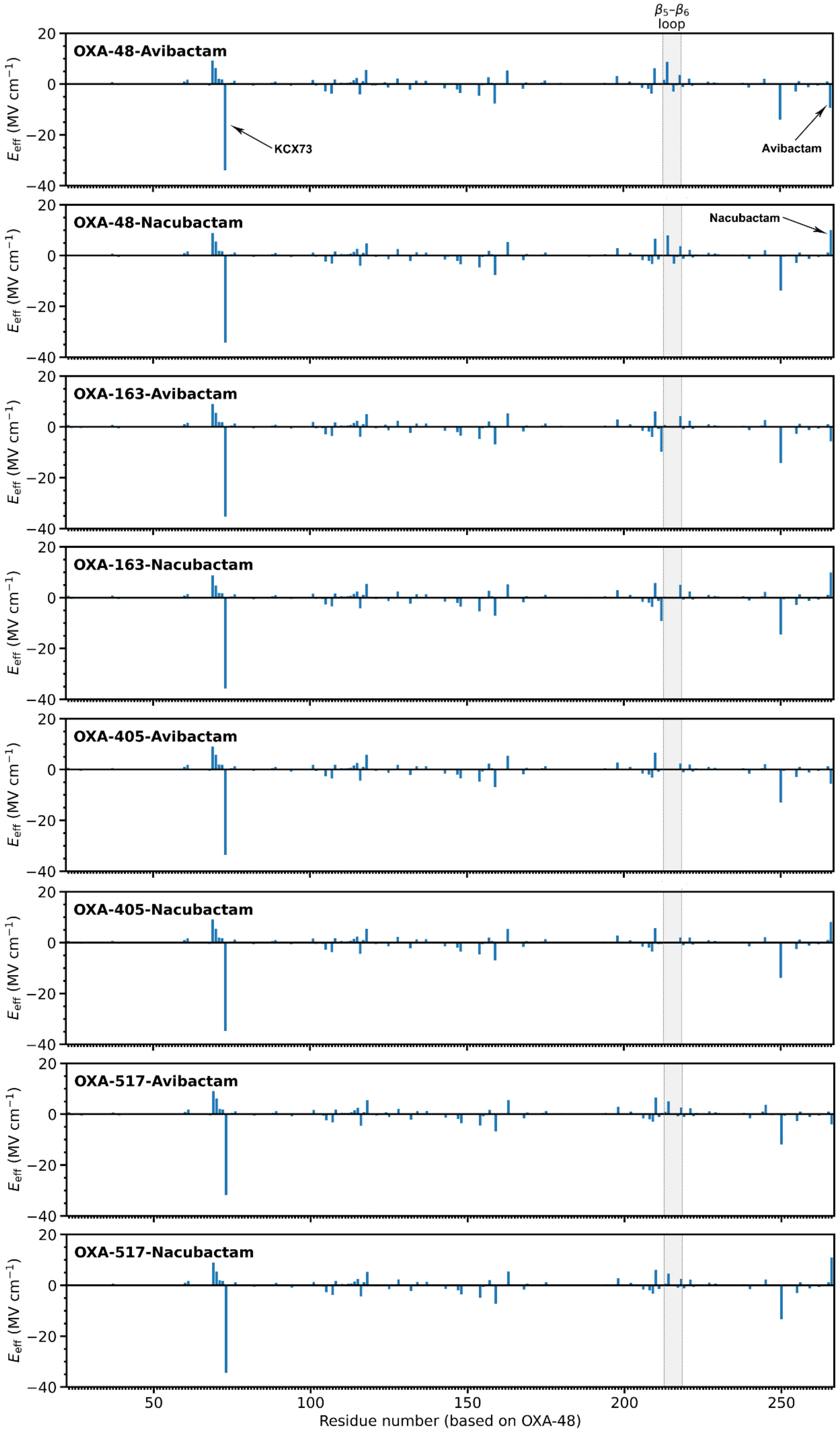

β5–β6
loop
OXA-48-Avibactam
KCX73
Avibactam
OXA-48-Nacubactam
Nacubactam
OXA-163-Avibactam
OXA-163-Nacubactam
OXA-405-Avibactam
OXA-405-Nacubactam
OXA-517-Avibactam
OXA-517-Nacubactam
$E_{eff}$ (MV cm$^{-1}$)
20
0
−20
−40
50
100
150
200
250
Residue number (based on OXA-48)

**Figure S12.** Effective electric field contributions of OXA-48 like enzymes and DBO projected along the DBO C-N bond, calculated using the transition state structures. The contributions from the carboxylated Lys73 (KCX) and DBOs are indicated by arrows. The negative electric field generated by KCX is overestimated because the proton associated with KCX is not included in the electric field analysis.

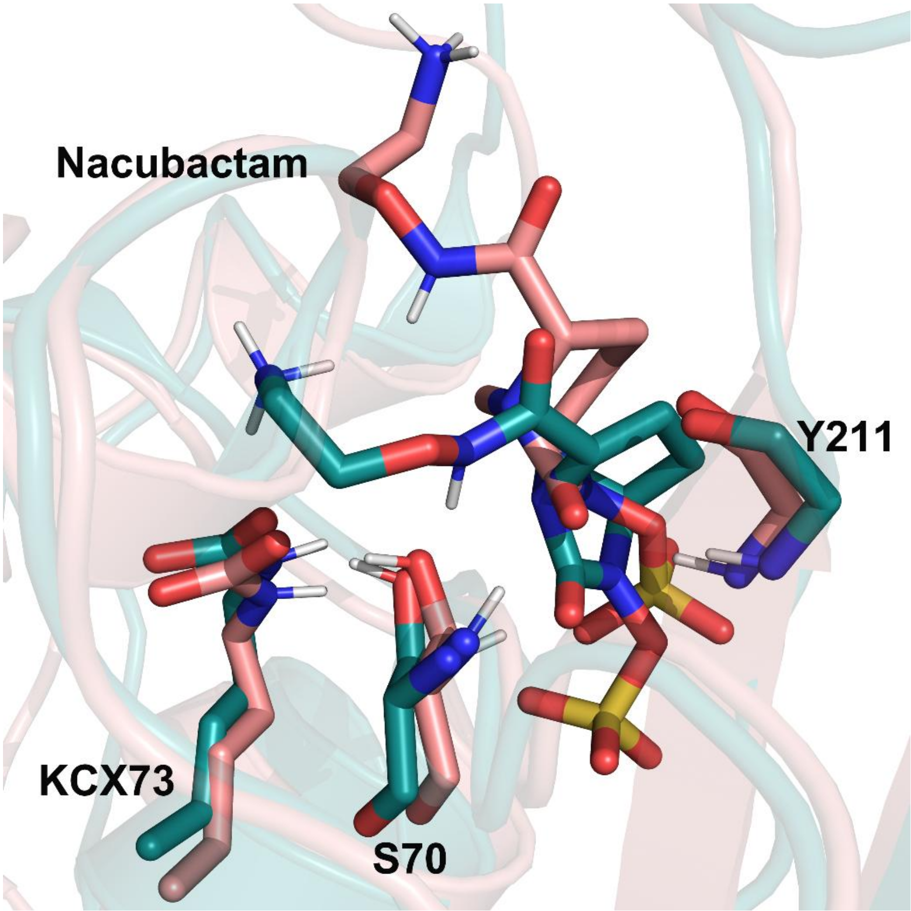


**Figure S13.** The nacubactam-bound complexes sample a distinct conformational space in relative binding free energy calculations (teal) compared with that in MM-PBSA binding energy calculations (pink). Final equilibrated structures of OXA-48 bound to nacubactam were aligned and are shown. The sidechain of carboxylated Lys73 (KCX), the backbone of Tyr211 and Ser70 and nacubactam are shown as sticks. Only polar hydrogens are shown for clarity. Nacubactam has 8.0, 7.8 and 9.2 kcal $mol^{-1}$ higher binding free energy than avibactam for OXA-48, -163, and -405, respectively (see also Table S5).

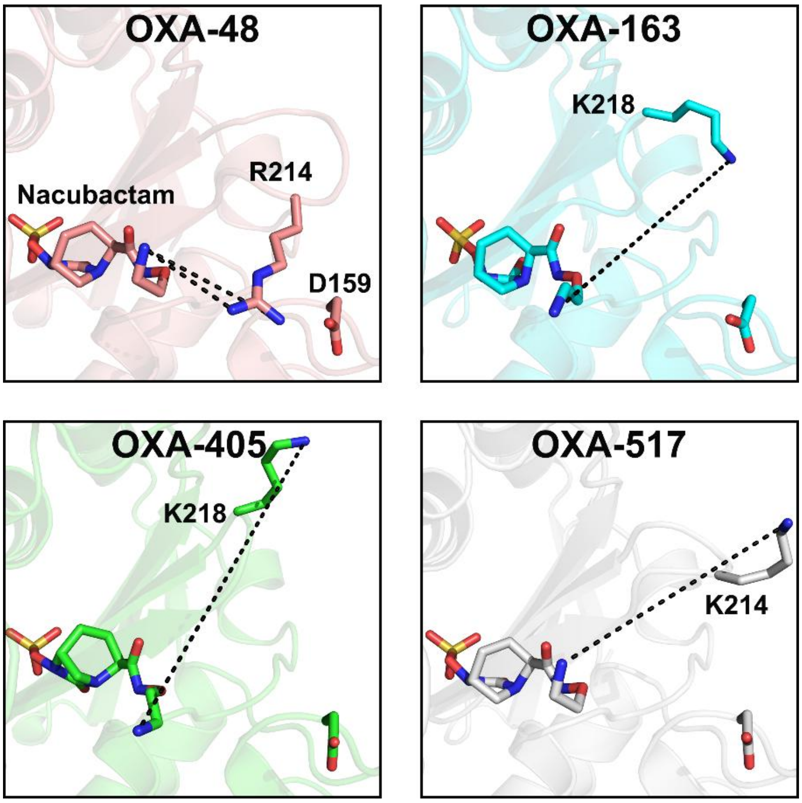


**Figure S14.** Atom-atom distances monitored (black dashed lines) for determining the closest distance between nacubactam charged nitrogen and Arg214 in OXA-48, Lys218 in OXA-163 and -405 or Lys214 in OXA-517 charged nitrogen during the 10 ns Michaelis complex MM MD simulations (see Figure S15). The crystal structures of nacubactam-bound OXA-48 (PDB ID: 9H12 [14], chain A), OXA-163 (PDB ID: 9HPV [14], chain A), OXA-405 (PDB ID: 9H18 [14], chain A) and the apo form of OXA-517 (PDB ID: 6HB8 [15], chain A) with modelled nacubactam are shown, with nacubactam and key residues displayed as sticks.

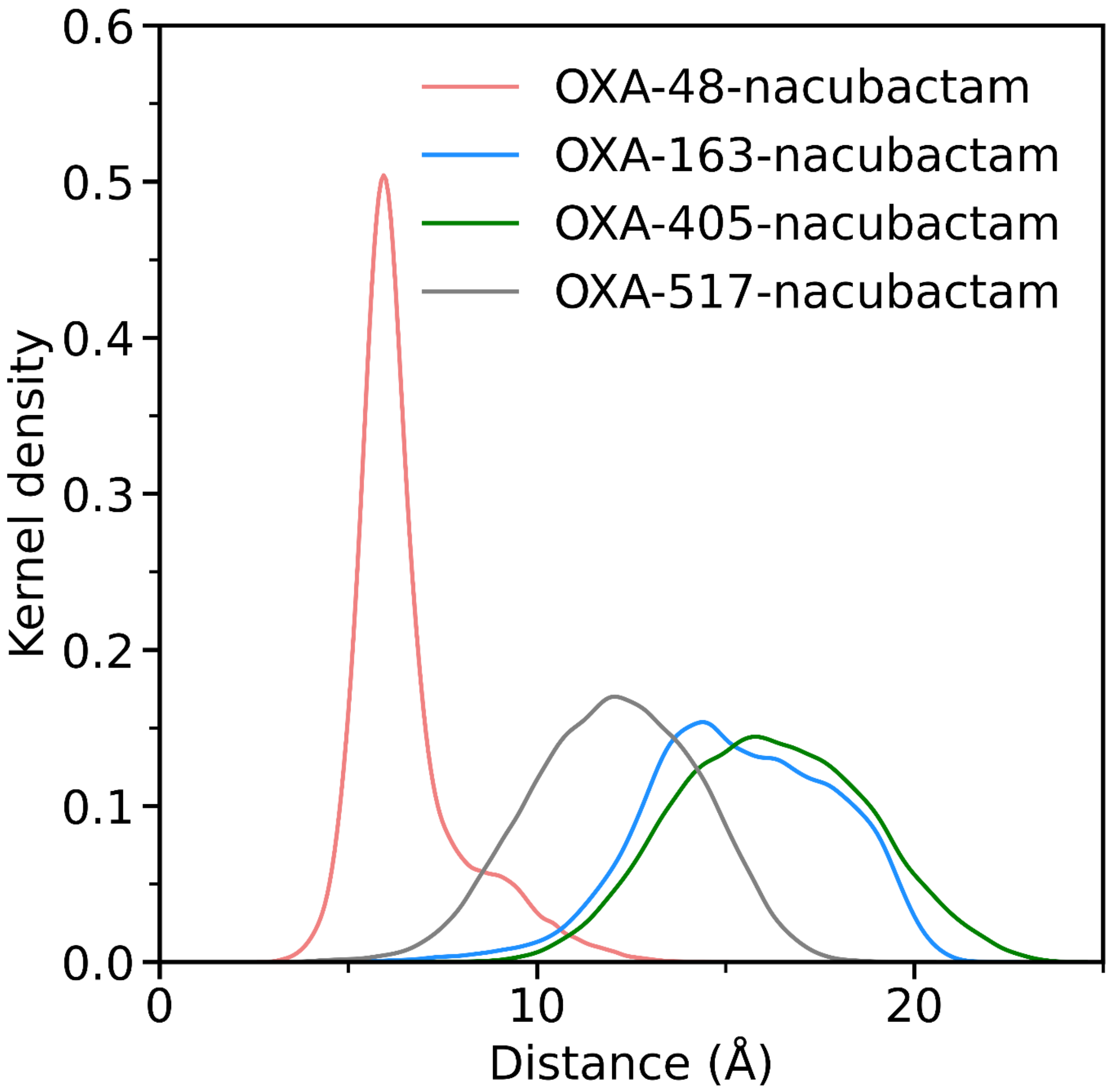


**Figure S15.** Distributions (kernel density estimates) of the closest distance between nacubactam ammonium group and Arg214 in OXA-48, Lys218 in OXA-163 and -405 or Lys214 in OXA-517 charged nitrogen during the 100 ns Michaelis complex MM MD simulations. The detailed distance measurements can be found in Figure S14.

**Table S1. Fraction of different hydration states observed around the DBO moiety during the 100 ns Michaelis complex MM MD simulations**

| Enzyme | Inhibitor | Hydration state 1 (with additional water) | Hydration state II (no additional water) |
|---|---|---|---|
| OXA-48 | Avibactam | 32.5% | 67.5% |
| | Nacubactam | 34.3% | 65.7% |
| OXA-163 | Avibactam | 27.3% | 72.7% |
| | Nacubactam | 24.6% | 75.4% |
| OXA-405 | Avibactam | 30.8% | 69.2% |
| | Nacubactam | 15.5% | 84.5% |
| OXA-517 | Avibactam | 36.1% | 63.9% |
| | Nacubactam | 35.7% | 64.3% |

**Table S2. Free energy barriers and average number of solvent H-bond to the DBO carbonyl oxygen during the second step of acylation of OXA-48 by avibactam and nacubactam from 2D QM/MM US MD simulations**

| Inhibitor | Replica number | Free energy barrier (kcal $mol^{-1}$) | Average solvent H-bond to the DBO carbonyl oxygen[a] |
|---|---|---|---|
| Avibactam | 1 | 10.8 | 0.01 |
| | 2 | 11.1 | 0.00 |
| | 3 | 8.7 | 0.00 |
| | 4 | 11.1 | < 0.01 |
| | 5 | 11.3 | 0.16 |
| Nacubactam | 1 | 12.4 | 0.28 |
| | 2 | 12.9 | 0.00 |
| | 3 | 12.9 | 0.00 |
| | 4 | 12.3 | 0.06 |
| | 5 | 13.2 | 0.07 |

[a] The average number of solvent H-bond was calculated based on the minimum free energy path trajectories.

**Table S3. Calculated free energy barriers for DBO acylation of OXA-48 and OXA-163**

| Enzyme | Inhibitor | Boltzmann-weighted first step free energy barrier (kcal mol$^{-1}$)[a] | Boltzmann-weighted second step free energy barrier (kcal mol$^{-1}$)[a] | Boltzmann-weighted overall free energy barrier (kcal mol$^{-1}$)[b] |
|---|---|---|---|---|
| OXA-48 | Avibactam | 12.9 ± 0.4 | 8.9 ± 0.6 | 12.4 ± 0.7 |
| | Nacubactam | 13.4 ± 0.5 | 12.6 ± 0.3 | 16.4 ± 0.5 |
| OXA-163 | Avibactam | 14.7 ± 0.6 | 7.8 ± 0.4 | 13.4 ± 0.7 |
| | Nacubactam | 14.3 ± 0.6 | 11.8 ± 0.5 | 17.8 ± 0.7 |

[a] The Boltzmann-weighted free energy barriers and Boltzmann-weighted standard deviations were obtained from five independent umbrella sampling calculations.

[b] The overall free energy barriers were obtained by adding two Boltzmann-weighted contributions: the free energy differences between the Michaelis complex and the tetrahedral intermediate ($\Delta G_1$), and the free energy barriers of the second step (from Table S3) ($\Delta^{\ddagger}G_2$). The standard deviations were calculated using the standard law of propagation of uncertainties ($\sigma = \sqrt{\sigma_{\Delta G1}{}^2 + \sigma_{\Delta\ddagger G2}{}^2}$).

**Table S4. Calculated free energy barriers for the second step of DBO acylation of OXA-405 and OXA-517**

| Enzyme | Inhibitor | Boltzmann-weighted second step free energy barrier (kcal mol$^{-1}$)[a] |
|---|---|---|
| OXA-405 | Avibactam | 9.5 ± 0.3 |
| | Nacubactam | 11.8 ± 0.6 |
| OXA-517 | Avibactam | 8.1 ± 0.4 |
| | Nacubactam | 11.5 ± 0.4 |

[a] The Boltzmann-weighted free energy barriers and Boltzmann-weighted standard deviations were obtained from five independent umbrella sampling calculations.

**Table S5. Calculated binding energies and relative binding free energies of DBO Michaelis complexes**

| Enzyme | Inhibitor | MM-PBSA binding energy $\Delta E$ (kcal mol$^{-1}$)[a] | MM-PBSA binding energy difference $\Delta\Delta E$ (kcal mol$^{-1}$)[b] | Relative binding free energy (kcal mol$^{-1}$)[c] |
|---|---|---|---|---|
| OXA-48 | Avibactam | −12.7 ± 1.0 | −7.9 ± 2.0 | −8.0 |
| | Nacubactam | −4.8 ± 1.7 | | |
| OXA-163 | Avibactam | −11.9 ± 0.9 | −5.2 ± 2.2 | −7.8 |
| | Nacubactam | −6.7 ± 2.0 | | |
| OXA-405 | Avibactam | −11.6 ± 0.7 | −5.5 ± 2.1 | −9.2 |
| | Nacubactam | −6.1 ± 2.0 | | |
| OXA-517 | Avibactam | −12.4 ± 0.8 | −7.2 ± 2.1 | ND |
| | Nacubactam | −5.2 ± 1.9 | | |

[a] Standard deviations obtained from MM-PBSA calculations on ten independent 10 ns MD trajectories.

[b] The binding energy differences were calculated with the MM-PBSA binding energy of avibactam referenced to that of nacubactam. The standard deviations were calculated using the standard law of propagation of uncertainties ($\sigma = \sqrt{\sigma_{\text{avibactam}}^2 + \sigma_{\text{nacubactam}}^2}$).

[c] Relative binding free energies were calculated using the alchemical transfer method with avibactam referenced to nacubactam (see Note S4). Single replicas were used, hence no standard deviation reported. ND = not determined.

**Table S6. Calculated $\log_{10}(k_2/K_i)$ derived from umbrella sampling reaction barriers and MM-PBSA binding energies for the acylation of OXA-48-like enzymes by DBOs**

| Enzyme | Inhibitor | $pIC_{50}$[a] | $\text{Log}_{10}(k_2/K_i)_{\text{calc}}$[b] |
|---|---|---|---|
| OXA-48 | Avibactam | 6.6 | 15.5 |
| | Nacubactam | 4.7 | 7.1 |
| OXA-163 | Avibactam | 6.7 | 15.8 |
| | Nacubactam | 5.9 | 9.1 |
| OXA-405 | Avibactam | 6 | 14.3 |
| | Nacubactam | 5.0 | 8.7 |
| OXA-517 | Avibactam | ND | 15.9 |
| | Nacubactam | ND | 8.2 |

[a]Data derived from experimental $IC_{50}$ values of ref. [14].

[b] The $k_2$ values were derived from the reaction barriers reported in Tables S3 and S4 using transition state theory [16]; the $K_i$ values were derived from the MM-PBSA binding energies reported in Table S5 using the thermodynamic relation between the Gibbs free energy and the equilibrium constant for non-covalent binding, $\Delta G = -RT\ln K$. ND = not determined.

(12) Bannwarth, C.; Ehlert, S.; Grimme, S. GFN2-xTB-An accurate and broadly parametrized self-consistent tight-binding quantum chemical method with multipole electrostatics and

density-dependent dispersion contributions. *J Chem Theory Comput* **2019**, *15* (3), 1652-1671. DOI: 10.1021/acs.jctc.8b01176.

(13) Tan, Z. Q.; Gallicchio, E.; Lapelosa, M.; Levy, R. M. Theory of binless multi-state free energy estimation with applications to protein-ligand binding. *J Chem Phys* **2012**, *136* (14), 144102. DOI: 10.1063/1.3701175.

(14) Hoff, J. F.; Goudar, K. E.; Calvopiña, K.; Beer, M.; Hinchliffe, P.; Shaw, J. M.; Tooke, C. L.; Takebayashi, Y.; Cadzow, A. F.; Harmer, N. J.; et al. Electrostatic interactions influence diazabicyclooctane inhibitor potency against OXA-48-like *β*-lactamases. *RSC Med Chem* **2025**, *16* (11), 5441-5455. DOI: 10.1039/d5md00512d.

(15) Dabos, L.; Raczynska, J. E.; Bogaerts, P.; Zavala, A.; Girlich, D.; Bonnin, R. A.; Dortet, L.; Peyrat, A.; Retailleau, P.; Iorga, B. I.; et al. Structural and biochemical features of OXA-517: a carbapenem and expanded-spectrum cephalosporin hydrolyzing OXA-48 variant. *Antimicrob Agents Chemother* **2023**, *67* (2), e0109522. DOI: 10.1128/aac.01095-22.

(16) Garcia-Viloca, M.; Gao, J.; Karplus, M.; Truhlar, D. G. How enzymes work: analysis by modern rate theory and computer simulations. *Science* **2004**, *303* (5655), 186-195. DOI: 10.1126/science.1088172.
